# BBSome-Mediated Clearance of Ubiquitinated IMPG2 Defines a Constitutive Ciliary Retrieval Pathway in Photoreceptors

**DOI:** 10.1101/2025.07.29.667331

**Authors:** Tirthasree Das, Gary A. Bradshaw, Jeanette Hyer, Markus Masek, Yien-Ming Kuo, Ruxandra Bachmann-Gagescu, Marian Kalocsay, Maxence V. Nachury

## Abstract

The BBSome mediates the retrieval of ubiquitinated membrane proteins from cilia, but its physiological cargoes in photoreceptors remain largely unidentified. Here, we find that K63-linked ubiquitin (UbK63) chains accumulate in the outer segment (OS, equivalent of cilia) of *Bbs4^−/−^* photoreceptors from the onset of OS formation. Through quantitative profiling of the UbK63-associated OS proteome, we identify the transmembrane fragment of interphotoreceptor matrix proteoglycan 2 (IMPG2^m^) as a principal cargo of the BBSome. In *Bbs4^−/−^* mice, ubiquitinated IMPG2^m^ aberrantly accumulates in OSs, and disruption of IMPG2^m^ ubiquitination impairs its retrieval and clearance. Because full-length IMPG2 traffics to the OS to deliver its extracellular domain to the matrix, our data support a model in which IMPG2^m^ undergoes constitutive cycling between the inner and outer segments. These findings redefine the BBSome’s role in photoreceptors from quality control to constitutive membrane protein turnover.

## INTRODUCTION

Bardet-Biedl Syndrome (BBS; OMIM 209900) is a Mendelian disorder characterized by retinal degeneration, polydactyly, obesity and kidney malformations, caused by mutations in any of at least 25 *BBS* genes^1^. Despite significant advances in understanding the molecular basis of BBS, no treatment currently exists for its retinal degeneration. The genetic complexity of BBS was considerably simplified by the discovery that eight BBS gene products form a stable complex, named the BBSome^2^, while most other BBS proteins function in the assembly or regulation of that complex^3,4^.

Studies in cultured mammalian cells and in the model unicellular organism *Chlamydomonas reinhardtii* have established that the BBSome retrieves membrane-associated proteins from cilia back to the cell body^5–7^. In mammals, the best characterized BBSome cargoes are G-protein-coupled receptors (GPCRs) that undergo regulated exit based on their activation state. In recent years, ubiquitination has emerged as a critical regulatory mechanism for ciliary GPCR retrieval. Long known for its role in tagging proteins for degradation or other regulatory processes^8,9^, ubiquitin (Ub) was recently shown to mark ciliary GPCRs for BBSome-mediated retrieval^10,11^. Polyubiquitin chain linkage can determine biological outcomes: lysine 48–linked ubiquitin (UbK48) chains typically drive proteasomal degradation of soluble proteins, whereas lysine 63–linked ubiquitin (UbK63) chains target membrane proteins for lysosomal degradation^8,9,12^ and mark ciliary GPCRs for retrieval^13^. Once ubiquitinated, ciliary GPCRs are recognized by the UbK63 chain reader TOM1L2, which latches ubiquitinated cargoes onto the BBSome^14^.

Whether this GPCR-based model extends to photoreceptors remains unclear, as rhodopsin—the principal photoreceptor GPCR—does not undergo regulated ciliary exit. Instead, some proteins such as the SNARE syntaxin-3, accumulate in photoreceptor outer segments (OSs; the equivalent of cilia) when BBSome function is compromised^15^. Given that rhodopsin’s tremendous membrane trafficking flux to the OS can drag along ‘bystander’ membrane proteins^16^, it has been proposed that such proteins, including syntaxin-3, accidentally enter the outer segment and are then retrieved by the BBSome to the inner segment (IS)^17^.

Additional support for the idea that membrane proteins can ‘leak’ from photoreceptor inner segments (ISs; equivalent of the soma) into outer segments and require retrieval has emerged from studies of transition zone (TZ) mutants. The TZ acts as a gate at the base of the cilium, preventing the free diffusion of membrane proteins between ciliary and plasma membranes, and consists of over 30 structural proteins such as CEP290, NPHP1, TCTN2^18,19^. Mice harboring mutations in these proteins exhibit phenotypes strikingly similar to those in *Bbs* mutants, including the inappropriate accumulation of select membrane proteins in the OS^20–22^. Based on these observations, a prevailing model proposes that the BBSome compensates for TZ-crossing accidents by retrieving proteins that erroneously enter the OS. Although compelling, this model does not explain why only certain proteins accumulate in BBSome-deficient OSs, nor does it clarify how the BBSome specifically recognizes these ‘accidental’ OS entrants.

Here, we show that K63-linked ubiquitinated proteins accumulate in the OS from the earliest stages of OS development in *Bbs4^−/−^* mutant mice. Purification of K63-linked ubiquitinated proteins from *Bbs4^−/−^* OSs identified the photoreceptor matrix transmembrane protein IMPG2^m^, and we show that ubiquitinated IMPG2^m^ accumulates in OS from *Bbs4^−/−^* mice. Further, mutation of the cytoplasmic lysines in IMPG2 blocks its retrieval from the OS to the IS. Because IMPG2^m^ helps deposit the photoreceptor matrix surrounding the OS, our data support a model in which IMPG2^m^ normally cycles into and out of the OS as part of normal photoreceptor biology, rather than merely gaining entry via gating errors subsequently corrected by a BBSome-mediated quality control mechanism.

## RESULTS

### K63-linked ubiquitin chains accumulate in photoreceptor outer segments in the absence of BBSome function

In past work, we observed a marked accumulation of ubiquitin in the OSs of *Bbs4^−/−^* mouse photoreceptors relative to WT controls at post-natal day 21 (P21)^10^. Because the morphology of photoreceptor OSs deteriorates when BBSome function is compromised^15,23–26^, we considered that ubiquitinated proteins might amass as a secondary consequence of OS degeneration. To test this hypothesis, we examined the timing of ubiquitin chain accumulation in *Bbs4^−/−^* OSs at the earliest developmental stages.

Surprisingly, a robust ubiquitin staining was detectable in *Bbs4^−/−^* OSs at P8 –when OSs are first forming^27,28^– (**Fig. 1A**) and persisted at P11 and P15 as OSs matured. Quantitation of the ratio of ubiquitin signal in OS vs. IS confirmed that ubiquitin accumulation in OS of *Bbs4^−/−^* photoreceptors remained relatively constant from P8 onward (**Fig. 1B**). This early and stable accumulation of ubiquitin in *Bbs4^−/−^* OSs is at odds with IS proteins slowly leaking into the OS and progressively accumulating when BBSome function is compromised.

**Figure 1:**
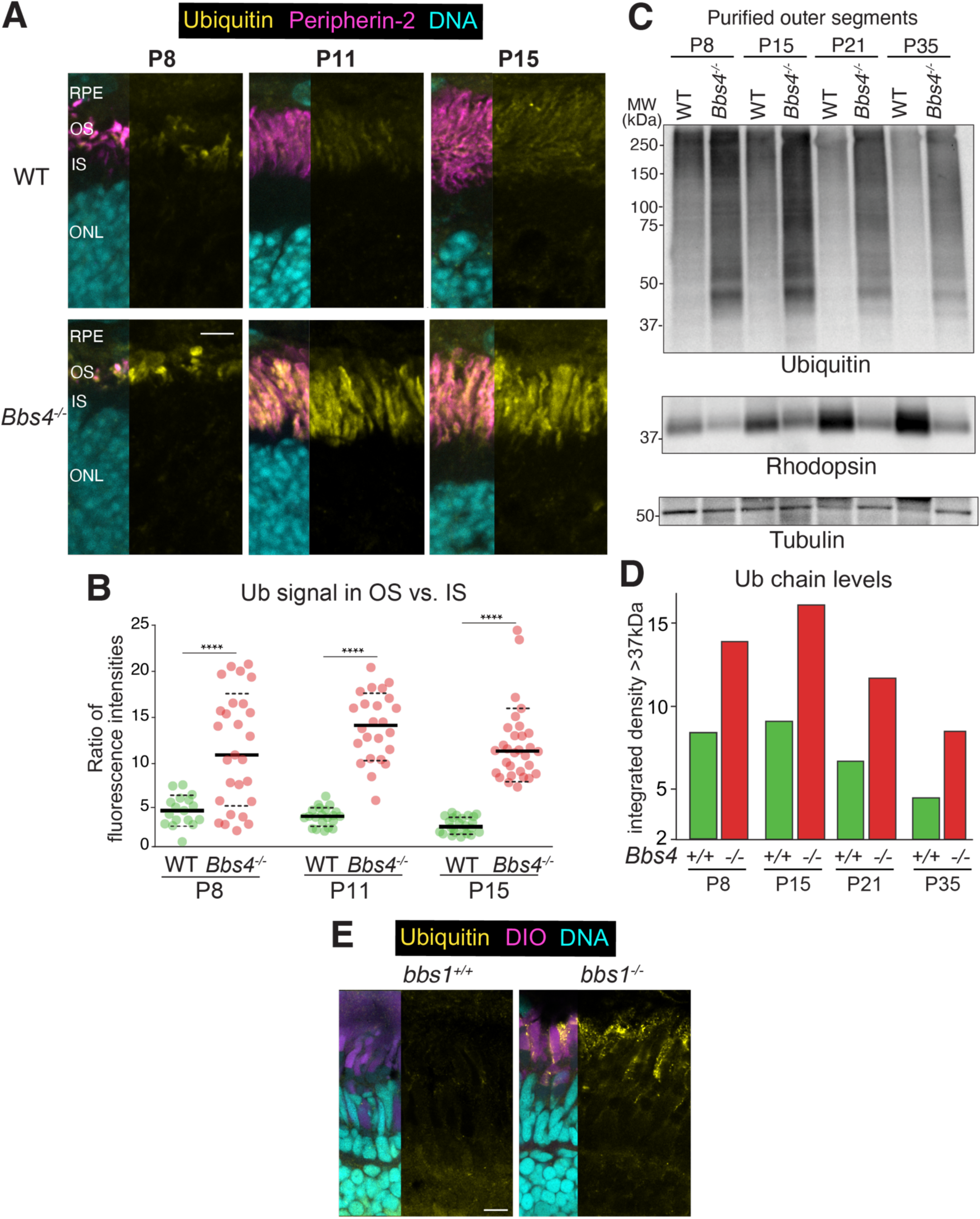
Ubiquitin accumulates in outer segments of BBSome-deficient photoreceptors as soon as photoreceptors form. **A-B.** Immunofluorescence analyses of mouse retina showing stable enrichment of ubiquitin in *Bbs4^−/−^* photoreceptor outer segments from P8 onwards. **A.** Representative images of retinal cryosections from *Bbs4^−/−^* and wild-type mice at post-natal day (P) 8, 11, and 15, stained for ubiquitin (FK2, yellow), peripherin-2 (magenta) and DNA (cyan). RPE: retinal pigmented epithelium, OS: outer segment, IS: inner segment, ONL: outer nuclear layer. Scale bar: 5μm. **B.** Quantitation of fluorescence signals for ubiquitin in the outer vs. inner segments at P8, P11 and P15. Ratios of outer to inner segment fluorescence intensities are plotted as dot plots (*N* = 2 retinas from 3 mice with 3 sections per animal for each genotype). Solid black lines represent medians; dashed lines represent the interquartile range. Statistical significance was determined by one-way ANOVA followed by Tukey’s post hoc test (****, *p* ≤ 0.0001). **C-D.** Biochemical analyses illustrating the time course of ubiquitin enrichment in *Bbs4^−/−^* photoreceptor outer segments. **C.** Outer segments from wild-type and *Bbs4^−/−^* retinas collected at the indicated ages were immunoblotted for ubiquitin (top), rhodopsin (middle) and tubulin (loading control, bottom). MW, molecular weight. **D.** Quantitation of the ubiquitin signal above 37 kDa from the blots shown in (**C**). Data are presented in a bar graph. **E.** Ubiquitin accumulation in zebrafish *bbs1^−/−^* photoreceptor outer segments. Representative images of zebrafish retinal sections from *bbs1^−/−^* and *bbs1^+/+^* (siblings of the mutants) animals at 34 days post-fertilization stained for ubiquitin (yellow), membranes (DiO, purple), and DNA (cyan). Scale bar: 10 μm.

Biochemical analysis of purified OS fractions from *Bbs4^−/−^* and WT mice further confirmed that ubiquitinated proteins – appearing as a high-molecular weight smear on Ub immunoblots– are enriched in *Bbs4^−/−^* OSs from P8 to P35 (**Fig. 1C-D**). The overall levels of ubiquitinated proteins in *Bbs4^−/−^* OSs remained relatively constant throughout photoreceptor development from their inception on, again suggesting that ubiquitinated proteins accumulate in *Bbs4^−/−^* OSs not simply because of a slow leaking corrected by the BBSome.

We next asked whether ubiquitin chains also accumulate in cone photoreceptors when BBSome function is compromised. Given that the mouse retina predominantly comprises rod photoreceptors, and that cone photoreceptors degenerate rapidly in *Bbs* knockout mice^26^, we turned our attention to the cone-dominant zebrafish model. Thanks to photoreceptor regeneration in zebrafish, *bbs1^−/−^* mutants retain cones as they age^29^. We found that *bbs1^−/−^* zebrafish displayed abundant ubiquitin staining in photoreceptor OSs, paralleling our findings in *Bbs4^−/−^* mouse rods (**Fig. 1E**).

Because previous studies in cultured kidney cells demonstrated that K63-linked (but not K48-linked) ubiquitin chains accumulate in cilia when BBSome function is lost^10,11^, we turned our attention to ubiquitin linkages. Consistent with these results, we detected no difference in K48- or K11-linked ubiquitin staining between wildtype and *Bbs4^−/−^* mouse photoreceptors (**Fig. S1A-B**). To detect K63 ubiquitin linkages, we employed Tandem Ubiquitin Binding Entities (TUBEs), which are proteins engineered to bind ubiquitin chains with high affinity and selectivity^30^. A TUBE that recognizes all Ub chains (pan-TUBE, based on HR23a^30^) replicated the accumulation of ubiquitin in *Bbs4^−/−^* OSs (**Fig. S1C**). A K63TUBE, which recognizes K63-linked ubiquitin (UbK63) chains with >1,000x selectivity over other chains^31^, revealed a similarly high and sustained enrichment of UbK63 chains in *Bbs4^−/−^* OSs from P8 onwards (**Fig. 2A-B**).

**Figure 2:**
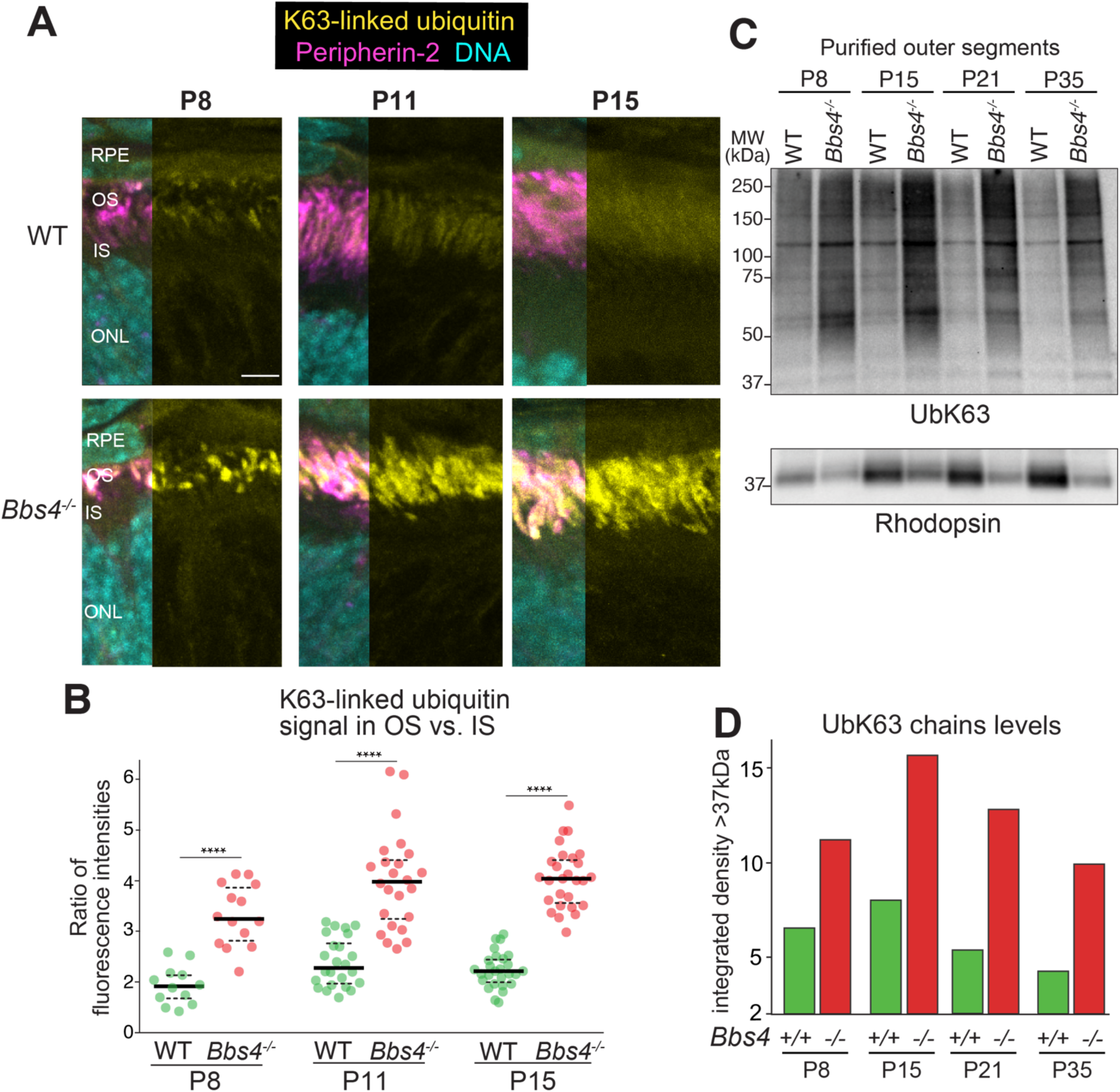
K63-linked ubiquitin chains accumulate in outer segments of *Bbs4^−/−^* photoreceptors as soon as photoreceptors form. **A-B**. Immunofluorescence analyses of mouse retina showing early and sustained enrichment of K63-linked ubiquitin chains in *Bbs4^−/−^* photoreceptor outer segments. **A.** Representative images of retinal cryosections from *Bbs4^−/−^* and wild-type mice at P8, P11 and P15 stained for K63-linked ubiquitin chains (UbK63, GST-K63TUBE, yellow), peripherin-2 (magenta) and DNA (cyan). RPE: retinal pigmented epithelium, OS: outer segment, IS: inner segment, ONL: outer nuclear layer. Scale bar: 5 μm. See **Fig S1E** for the GST-only control. **B.** Quantitation of fluorescence signals for K63-linked ubiquitin chains in the outer vs. inner segments at P8, P11 and P15. Ratios of outer to inner segment fluorescence intensities are plotted as dot plots (*N* = 3 mice for each genotype, each with 2 retinas and 3 sections per retina analyzed). Solid black lines represent medians; dashed lines represent the interquartile range. Statistical significance was determined by one-way ANOVA followed by Tukey’s post hoc test (****, *p* ≤ 0.0001). **C-D.** Biochemical analyses illustrating the time course of K63-linked ubiquitin chains enrichment in *Bbs4^−/−^* photoreceptor outer segments. **C.** Outer segments isolated from wild-type and *Bbs4^−/−^* retinas at the indicated ages were immunoblotted for UbK63 (Apu3, top) and rhodopsin (1D4, bottom). **D.** Quantitation of the UbK63 signal above 37 kDa from the blots shown in (**C**). Data are presented in a bar graph.

Immunoblotting purified OSs fractions from P8 to P35 with a UbK63 linkage-specific antibody recapitulated these data, with pronounced UbK63 chain accumulation throughout development of *Bbs4^−/−^* OSs (**Fig. 2C-D**). Thus, rather than gradually accumulating with age, K63-linked ubiquitinated proteins rapidly reach –and maintain– a steady-state level in *Bbs4^−/−^* OSs at the onset of OS biogenesis.

Taken together, our data show that the accumulation of ubiquitin –particularly K63-linked ubiquitin chains– upon loss of BBSome function extends from primary cilia of cultured kidney cells to the specialized cilia of rod and cone photoreceptors, as well as to *Chlamydomonas* flagella. This strongly suggests that the BBSome has a conserved role in ferrying K63-linked ubiquitinated proteins out of cilia.

### The BBSome-associated UbK63 reader TOM1L2 accumulates in outer segments when BBSome function is compromised

We recently discovered that the BBSome engages the endosomal UbK63 reader TOM1L2 to retrieve UbK63-modified GPCRs from cilia^14^. Notably, a proteomics analysis of proteins associated with the plasma membrane of photoreceptor OSs identified seven out of eight BBSome subunits, along with TOM1L2, among the 65 most enriched proteins^32^. This finding suggested that TOM1L2 may collaborate with the BBSome to retrieve UbK63-modified cargoes from OSs.

In support of this hypothesis, an antibody that recognizes TOM1L2 and its paralogue TOM1 revealed a markedly increased OS staining in *Bbs4^−/−^* photoreceptors (**Fig. S2A-B**). Despite total TOM1L2 levels remaining unchanged between WT and *Bbs4^−/−^* retina, isolated OSs from *Bbs4^−/−^* animals revealed a ∼4.5-fold increase in TOM1L2 levels relative to WT (**Fig. S2C-D**). Echoing our findings, prior proteomics profiling of zebrafish *bbs1^−/−^* OS detected a highly significant 1.8-fold increase in TOM1L2 levels in *bbs1^−/−^* OS relative to *bbs1^+/+^*^29^. These results suggest that TOM1L2 acts as a bridge between the BBSome and UbK63-modified cargoes in multiple cell types including kidney cells, RPE cells and photoreceptors.

### Selective capture of UbK63-associated proteins from *Bbs4^−/−^* outer segments

Studies in cultured cells have shown that UbK63 chains mark ciliary GPCRs for regulated retrieval. Yet, opsins, the principle GPCRs in photoreceptors, are not thought to undergo retrieval from OS to IS. This disconnect raised the question of which proteins in the OS are modified with UbK63 chains and destined for retrieval back to the IS. To address this question, we devised a biochemical strategy to capture UbK63 chains and associated proteins from purified OSs of *Bbs4^−/−^* photoreceptors (**Fig. 3A**). We leveraged the K63TUBE, which possesses a K_d_ of ∼5 nM for UbK63 chains and over 1,000 selectivity for K63 linkages over K48, K11, or M11 linkages^31^. As observed in other contexts^30^, adding GST-K63TUBE during the lysis of OS fractions protected Ub chains from endogenous deubiquitinases (**Fig. S3A**). Biochemical analysis of Ub (**Fig. S3A**) or UbK63 (**Fig. 3B**) chains revealed a ∼4-fold increase in *Bbs4^−/−^* OSs relative to WT, and a similar enrichment in K63TUBE eluates, confirming the reproducibility and efficiency of the K63TUBE capture method.

**Figure 3:**
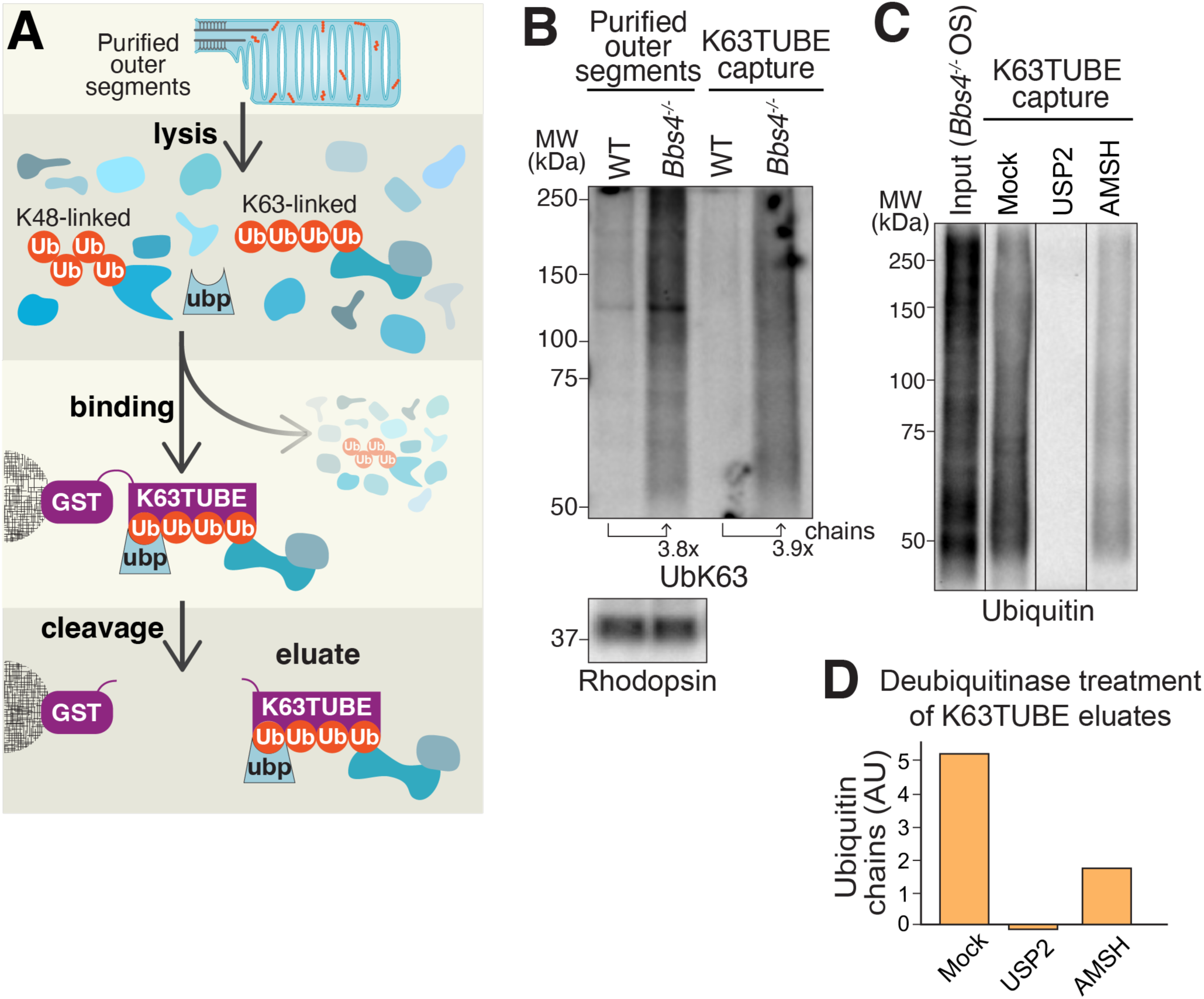
Biochemical isolation of K63-linked ubiquitinated proteins from photoreceptor outer segments. **A.** Diagram of the biochemical strategy. Recombinant GST-K63TUBE was incubated with lysates of purified OSs from wild-type or *Bbs4^−/−^* mice retinas. Proteins associated with UbK63 chains, covalently ubiquitinated proteins and ubiquitin-binding proteins (ubp), were captured on glutathione resin. Cleavage with HRV3C protease released K63TUBE and its bound proteins. **B.** K63TUBE faithfully captures K63Ub chains from photoreceptor OSs. Inputs (purified OSs mixed with K63TUBE) and K63TUBE eluates were probed for K63-linked ubiquitin (Apu3 antibody, top) or rhodopsin (loading control for inputs, bottom). 0.1% input and 20% of cleaved eluates were loaded for immunoblotting. The integrated intensity of the smear above 37 kDa normalized to WT and is shown below each lane. Purified OS lanes and K63TUBE capture lanes were quantified separately. See **Fig S3A** for additional experiment including a GST-only control. **C-D.** Deubiquitinase-based analysis of ubiquitin chain architecture. **C.** K63TUBE eluates from *Bbs4^−/−^* outer segment proteins were treated with AMSH (specific for the K63 linkage) or USP2 (cleaves all ubiquitin linkages) and probed for ubiquitin (FK2). **D.** The integrated intensity of the smear above 37 kDa from (**C**) is plotted in a bar graph. See **Fig S3B-C** for equivalent assays with WT outer segments.

We next evaluated the selectivity of UbK63 chain capture by treating the K63TUBE eluates with deubiquitinating enzymes (DUBs) with defined linkage specificity^33^. USP2, which cleaves all linkage types, removed all polyubiquitin signals (**Fig. 3C-D and S3B-C**). Meanwhile, AMSH, which specifically cleaves UbK63 linkages, reduced the polyubiquitin signal to about one-third of its abundance in mock-treated samples. These results indicate that most ubiquitin chains captured by the K63TUBE are K63-linked, thereby validating the specificity of our purification strategy.

### Identification of the K63Ub-associated proteome from photoreceptor outer segments

To identify the UbK63-marked proteins that accumulate in *Bbs4^−/−^* OSs, we performed a quantitative mass spectrometry analysis of the GST-K63TUBE eluates from WT and *Bbs4^−/−^* OSs (**Fig. 4A**). We leveraged tandem mass tag (TMT) multiplexed quantitative mass spectrometry for highly accurate relative protein quantitation across multiple samples^34–36^ and conducted the purifications in triplicate to allow for statistical analyses of relative enrichments between *Bbs4^−/−^* and WT samples. Strikingly, only five proteins met our significance criteria (*p* < 0.01, enrichment ≥1.75 fold; **Fig 4B**): ubiquitin, junctional adhesion molecule2 (JAM2), syntaxin-3 (STX3), the interphotoreceptor matrix proteoglycan 2 (IMPG2), and the amino acid transporter SLC3A2.

**Figure 4.**
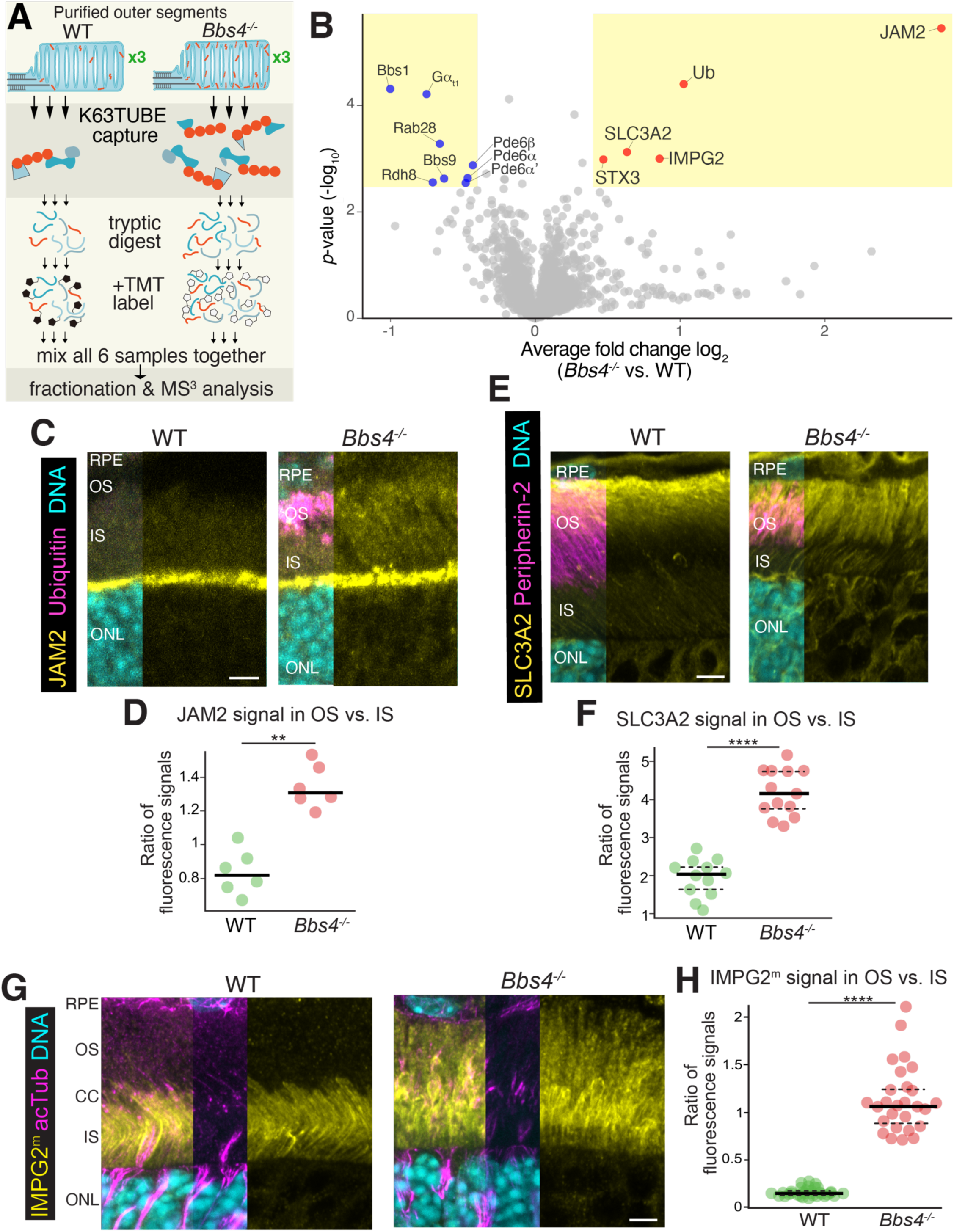
Proteomic profiling of the K63Ub-associated proteome from *Bbs4^−/−^* outer segments. **A.** Workflow of the proteomic analysis. OSs from wild-type and *Bbs4^−/−^* retinas (P15 animals) were prepared in biological triplicates and subjected to GST-K63TUBE capture. After on-bead digestion with trypsin, samples were labeled with a unique tandem mass tag (TMT), pooled, and peptides analyzed by synchronous precursor selection MS^3^ for both identification and quantitation. **B.** Volcano plot of statistical significance versus enrichment for *Bbs4^−/−^* compared to wild-type K63TUBE eluates. For 1,899 quantified proteins, statistical significance (-log10 *p* values of unpaired t-tests) is plotted against the TMT ratios of *Bbs4^−/−^*over WT samples. Proteins with a *p* value > 2.5 and a fold change > 0.3 are highlighted (yellow box). Significantly enriched proteins are shown in red and depleted proteins in blue. **C-D.** Accumulation of JAM2 in *Bbs4^−/−^* outer segments. **C.** Representative images of retinal sections (P15 animals) stained for JAM2 (yellow), ubiquitin (magenta) and DNA (cyan). Scale bar: 5 μm. **D.** Ratios of JAM2 fluorescence intensity in outer vs. inner segments. *N* = 1 retina from 2 mice, each with 1 section per animal for each genotype. Statistical significance was determined by an unpaired, non-parametric Mann-Whitney test (**, *p* < 0.005). **E-F.** Accumulation of SLC3A2 in *Bbs4^−/−^* outer segments. **E.** Representative images of retinal sections (P21) stained for SLC3A2 (yellow), peripherin-2 (magenta) and DNA (cyan). Scale bar: 5 μm. See **Fig S3D** for staining at P15. **F.** Ratios of SLC3A2 fluorescence intensity in outer vs. inner segments at P15. *N* = 2 retinas from 2 mice with 2 sections per animal for each genotype. Statistical significance was determined by an unpaired, non-parametric Mann-Whitney test (****, *p* ≤ 0.0001). **G-H.** Accumulation of IMPG2 in *Bbs4^−/−^* outer segments. **G.** Representative images of retinal sections (P15) stained for IMPG2 (yellow), acetylated tubulin (acTub, magenta) and DNA (cyan). Scale bar: 5 μm. **H.** Ratios of IMPG2 fluorescence intensity in outer vs. inner segments at P15. *N* = 2 retinas from 3 mice with 3 sections for each genotype. Statistical significance was determined by an unpaired, non-parametric Mann-Whitney test (****, *p* ≤ 0.0001).

The robust recovery of ubiquitin aligns with our immunofluorescence and immunoblotting results, providing internal validation for our workflow. Syntaxin-3, a SNARE protein involved in synaptic vesicle fusion^37^ and in the delivery of rhodopsin carrier vesicles to the OS^38,39^, typically localizes to the IS but abnormally accumulates in the OS of *Bbs* mice^15,40^. Our results suggest that it may be trafficked out of the OS via the BBSome and UbK63 chains. Alternatively, since the TUBE captures recover both ubiquitinated proteins and proteins associated with ubiquitin chains (e.g., TOM1L2, see Fig. **S2D**), syntaxin-3 may have been captured by the K63TUBE via non-covalent interactions with UbK63 chains due to its ability to directly bind to them^41^.

We next examined the remaining hits from the *Bbs4^−/−^* OS K63Ub-associated proteome by staining *Bbs4^−/−^* and WT retina. JAM2, a known tight junction component localized to the outer limiting membrane of photoreceptors^42^, showed a modest but reproducible and significant accumulation in OS *Bbs4^−/−^* OSs (**Fig. 4C-D**). SLC3A2, a solute carrier protein that forms heterodimeric amino acid transporters with members of the SLC7 family, is enriched in the plasma membrane of rod OS compared to disks^43^ and detected in OS-derived extracellular vesicles^44^. In WT retina, SLC3A2 localized predominantly in the distal region of the photoreceptor layer, indicative of a localization to RPE microvilli (**Fig. 4E-F, S3D**). In contrast, *Bbs4^−/−^* retina displayed uniform SLC3A2 labeling throughout the OS layer, suggesting that a fraction of SLC3A2 depends on the BBSome to traffic out of the OS.

Lastly, the interphotoreceptor matrix proteoglycans IMPG1 and IMPG2 are the chief constituents of the interphotoreceptor matrix^45^. IMPG1 is a secreted molecule, whereas IMPG2 is a transmembrane protein that sheds its large extracellular domain (IMPG2^ec^), which surrounds the photoreceptor OS and IS^46^. Autoproteolysis of IMPG2 produces IMPG2^ec^ and a transmembrane fragment (IMPG2^m^), both of which self-associate^46^. In WT photoreceptors, IMPG2^m^ predominantly resides in the IS^46,47^, whereas in *Bbs4^−/−^* retina IMPG2^m^ was present at similar levels in both IS and OS (**Fig. 4G-H**), suggesting that IMPG2^m^ separates from IMPG2^ec^ in the OS and is then marked with UbK63 chains for BBSome-mediated retrieval.

These combined findings confirm that each of the proteins in the OS K63Ub-associated proteome accumulates in the OS when BBSome function is disrupted, indicating that UbK63-dependent retrieval of these cargoes from the OS to the IS is impaired in *Bbs4*^−/−^ photoreceptors.

We note that eight proteins were significantly depleted in *Bbs4^−/−^* K63TUBE eluates from OSs relative to WT (**Fig. 4B**), including two subunits of the BBSome, Gα_t1_ transducin, the phosphodiesterase PDE6, RAB28 and the retinol dehydrogenase RDH8. The finding of BBSome subunits reduced in *Bbs4^−/−^* K63TUBE eluates relative to WT is consistent with findings that partially assembled BBSomes can fail to enter photoreceptor OSs^48^.

### Ubiquitinated IMPG2^m^ accumulates in *Bbs* outer segments

We next sought to determine whether proteins in the K63Ub-associated proteome of *Bbs* OSs are modified with ubiquitin chains. Since the TUBE strategy recovers both ubiquitinated proteins and proteins associated with ubiquitin chains (e.g. TOM1L2, see **Fig. S2D**), we switched to purifications under denaturing conditions to only capture proteins covalently linked to ubiquitin (**Fig. 5A**). OtUBD, a tightly folded ubiquitin binding domain with a remarkably high affinity for ubiquitin and resistance to denaturing agents^49^, readily captured ubiquitinated proteins from HEK cell lysates under denaturing conditions (**Fig. S4A**) and from OS extracts (**Fig. 5B** and **S4B-C**). Mirroring the K63TUBE capture, a broad ubiquitin smear was recovered by OtUBD from both WT and *Bbs4^−/−^* OSs, with a nearly two-fold increase in total Ub signal in *Bbs4^−/−^* compared to WT eluates.

**Figure 5:**
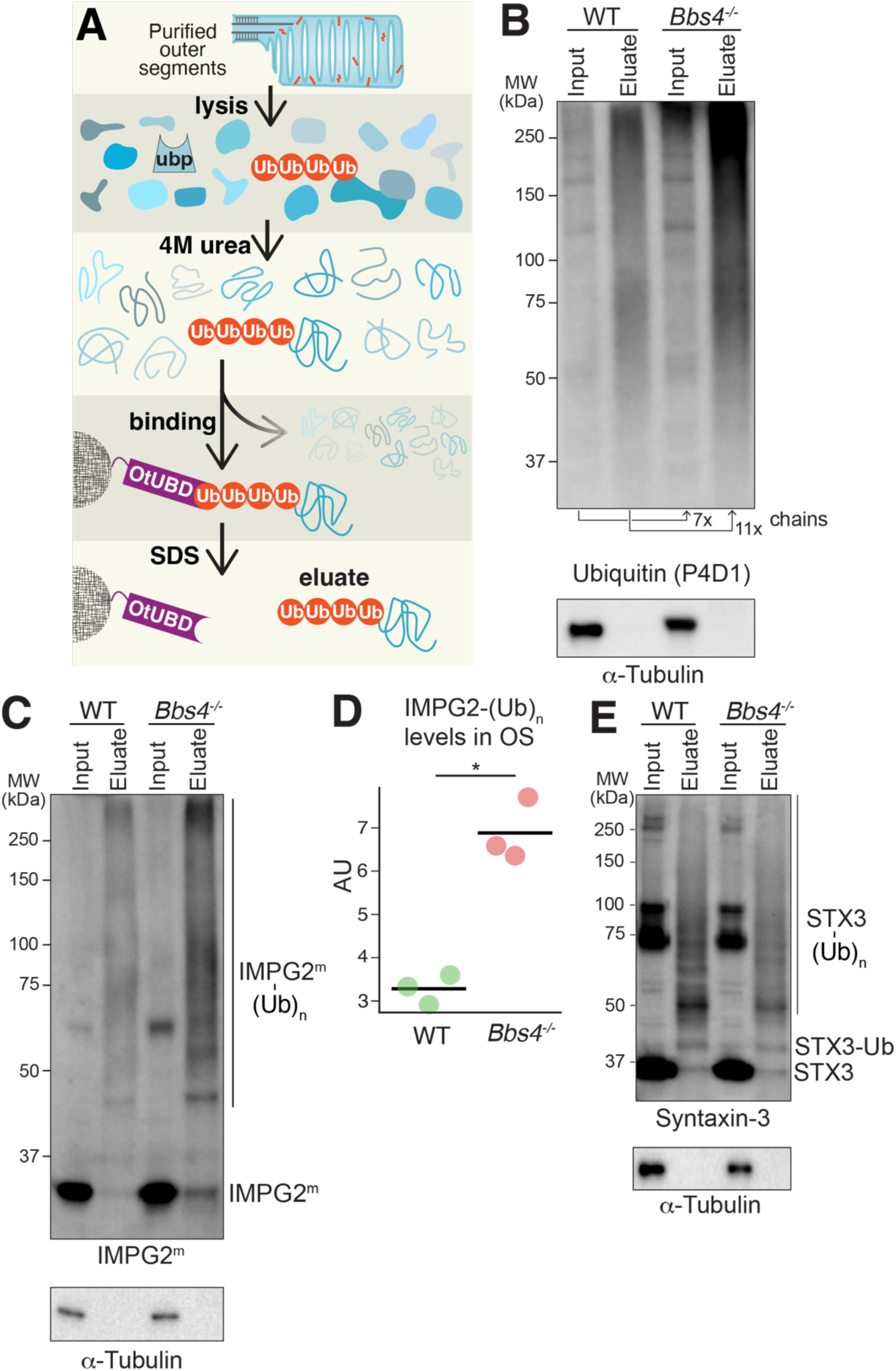
Ubiquitination of IMPG2^m^ is elevated in *Bbs4^−/−^* outer segments. **A.** Diagram of the biochemical strategy used to capture proteins covalently linked to ubiquitin. Purified OSs from WT and *Bbs4^−/−^* retinas (P15) were lysed and denaturated in 8 M urea, then diluted to 4 M urea to allow partial refolding of ubiquitin. Ubiquitinated proteins were subsequently captured on immobilized OtUBD and eluted by boiling in SDS sample buffer. **B.** Validation of the biochemical strategy. Purified OSs (input) and OtUBD eluates were probed for ubiquitin (P4D1, top panel) or tubulin (loading control, bottom panel). 2% input and 15% of eluates were loaded. The integrated intensity of the smear above 37 kDa was normalized to the WT value and is shown below each lane. **C-D.** Ubiquitination of IMPG2^m^ in *Bbs4^−/−^* and WT outer segments. **C.** Eluates from the OtUBD capture were probed for IMPG2^m^ (top panel) and tubulin (loading control, bottom panel). 2% input and 20% of eluates were loaded. **D.** The integrated intensity of the IMPG2^m^ smear above 37 kDa was measured (from **C**) and plotted as a dot plot. *N* = 3 animals per genotype. Statistical significance was determined by a paired parametric *t*-test (*, *p* < 0.05). **E.** Ubiquitination of syntaxin-3 (STX3) in *Bbs4^−/−^* and WT outer segments. Eluates from the OtUBD capture were probed for syntaxin-3 (top panel) and tubulin (loading control, bottom panel). 2% input and 25% of eluates were loaded.

Remarkably, OtUBD recovered very little IMPG2^m^ at the canonical molecular weight of 32 kDa (**Fig. 5C**). Instead, most of the IMPG2^m^ detected in OtUBD eluates migrated as high molecular weight bands indicative of mono- and poly-ubiquitinated species. Importantly, the signal of ubiquitinated IMPG2^m^ was measurably and significantly increased in OtUBD captures from *Bbs4^−/−^* OSs compared to WT OSs (**Fig. 5C-D**). This result indicates that ubiquitinated IMPG2^m^ accumulates in OS of *Bbs4^−/−^* photoreceptors, suggesting that ubiquitin chains are added onto IMPG2^m^ prior to BBSome-mediated transport back to the IS.

Similarly, most of syntaxin-3 in the OtUBD eluates migrated as high molecular weight species. However, the amount of ubiquitinated syntaxin-3 was not elevated in *Bbs4^−/−^* OSs compared to WT OSs (**Fig 5E**), suggesting that syntaxin-3 may not represents a BBSome cargo tagged with ubiquitin chains prior to BBSome-mediated exit from the OS.

### BBSome-dependent and ubiquitin-independent removal of syntaxin-3 from cilia

We next investigated whether ubiquitination and the BBSome participate in the exit of syntaxin-3 from cilia. Prior studies found that syntaxin-3 undergoes mono-ubiquitination in kidney cells and plays a role in exosome biogenesis^41^. Encouraged by the detection of syntaxin-3 in kidney cells^41^, we stained mouse kidney (IMCD3) cells for syntaxin-3. While syntaxin-3 was nearly undetectable in cilia of WT cells, deletion of the BBSome regulators *Arl6* or *Ift27* led to robust syntaxin-3 levels inside cilia (**Fig 6A-B**). The detection of syntaxin-3 inside IMCD3 cilia was specific, as siRNA-mediated depletion of syntaxin-3 reduced the ciliary signal to non-detectable levels (**Fig. S4D**). The accumulation of syntaxin-3 inside cilia when BBSome function is compromised mirrors similar results in photoreceptors, indicating a broad requirement for the BBSome in trafficking syntaxin-3 out of cilia and demonstrating that cultured cells can model the ciliary trafficking of syntaxin-3.

**Figure 6:**
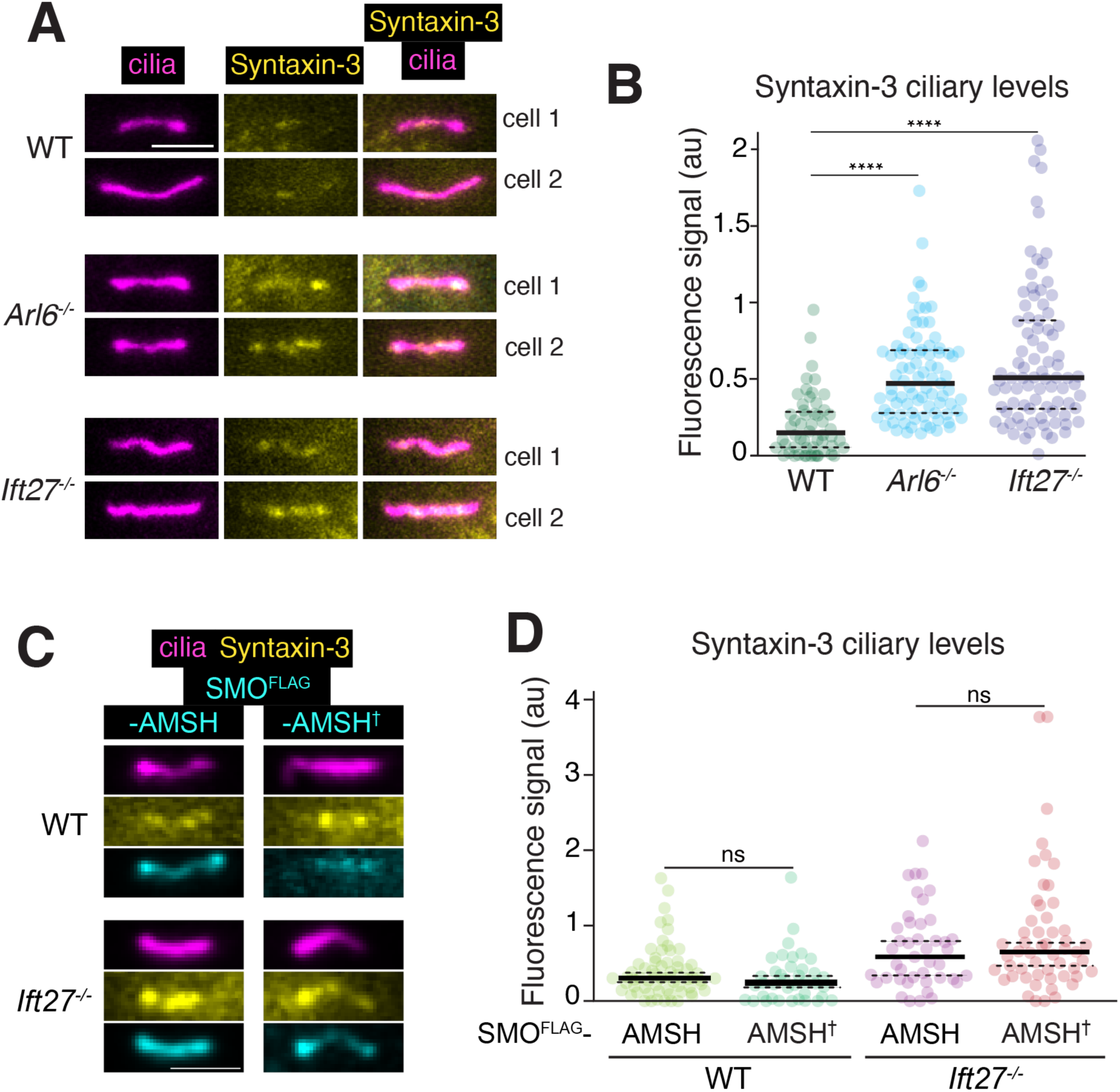
Syntaxin-3 accumulates in cilia of BBSome-deficient cells, but not in a UbK63-regulated manner. **A-B.** Accumulation of syntaxin-3 in primary cilia of BBSome-deficient cells. **A.** WT, *Arl6^−/−^*, and *Ift27^−/−^* IMCD3 cells were serum-starved, then fixed and stained for acetylated tubulin (cilia; magenta) and syntaxin-3 (yellow). Scale bar: 2μm. **B.** The fluorescence intensity of syntaxin-3 in cilia was quantified in each cell line, and plotted as dot plots. Statistical significance was determined by one-way ANOVA with Tukey’s post hoc test (****, *p* < 0.0001, *N ≥* 60 cilia). **C-D.** Removal of K63Ub chains from cilia does not affect syntaxin-3 accumulation. **C.** Representative images of cilia from WT and *Ift27^−/−^* IMCD3 cells expressing FLAG-tagged SMO (SMO^FLAG^) fused to AMSH (K63-specific deubiquitinase) or its catalytically dead variant (AMSH^✝^). Cells were transfected, serum-starved and stained for syntaxin-3 (yellow), acetylated tubulin (cilia; magenta), and FLAG (cyan). Channels are shifted to facilitate visualization. Scale bar: 2 μm. **D.** The ciliary fluorescence intensity of syntaxin-3 was measured under each condition and plotted as dot plots. Statistical significance was determined by one-way ANOVA with Tukey’s post hoc test (ns, not significant).

We leveraged the IMCD3 cell culture system to test whether ubiquitination gates the removal of syntaxin-3 from cilia. Previous work has shown that cleavage of K63-linked Ub chains inside cilia by the deubiquitinase AMSH results in the accumulation of BBSome cargoes inside cilia^10,50^. Congruently, stably expressed AMSH fused to the ciliary signaling receptor Smoothened (SMO-AMSH) accumulated to high levels in cilia. However, syntaxin-3 did not accumulate in cilia of SMO-AMSH cells compared to cells expressing a catalytically dead variant (SMO-AMSH^†^) (**Fig 6C-D**). We then considered that syntaxin-3 may accumulate in cilia when BBSome function fails because it recognizes UbK63 chains^41^ and UbK63 chains accumulate in *Bbs* cilia^10^. However, in *Ift27^−/−^* cells where the ciliary levels of syntaxin-3 are elevated compared to WT cells, removal of UbK63 chains from cilia did not alter the ciliary levels of syntaxin-3 (**Fig 6C-D**). These results demonstrate that ciliary K63-linked ubiquitination neither traps syntaxin-3 inside cilia nor promotes the exit of syntaxin-3 from cilia. These results contrast with previously characterized BBSome cargoes in cultured cells, which are fully reliant on K63Ub chains for removal from cilia^10,11^. Considering that syntaxin-3 ubiquitination is not increased in *Bbs* OSs compared to WT OSs, our data suggest that syntaxin-3 is not a conventional BBSome cargo. Instead, the accumulation of syntaxin-3 inside *Bbs* cilia and OSs may be indirectly caused by retrieval defects.

### Ubiquitination of Impg2^m^ is required for retrieval from photoreceptor outer segments

The OS K63Ub-associated proteome and subsequent biochemical validations demonstrate that IMPG2^m^ is modified with K63Ub chains in photoreceptor OSs. Additionally, IMPG2^m^ accumulates in the OS of *Bbs4^−/−^* photoreceptors. These data suggest that after escorting IMPG2^ec^ into the OS, IMPG2^m^ becomes marked with K63Ub chains, committing this protein for BBSome-mediated retrieval back to the IS.

To test the importance of ubiquitination in IMPG2^m^ trafficking from OS back to IS, we substituted all four lysine residues within the cytoplasmic tail of IMPG2 with arginine to generate IMPG2[cK0], a variant refractory to ubiquitination. Photoreceptor-selective expression of full-length IMPG2 was driven by the rhodopsin kinase promoter and gene delivery was carried out by packaging into adenovirus-associated virus AAV5 and injection into the mouse subretinal space (**Fig. 7A**). Expression of WT IMPG2 yielded a modest increase in the IMPG2^m^ signal in the IS, with no signal detectable in the OS (**Fig. 7B**). Thus, mild overexpression of IMPG2 does not perturb its trafficking in photoreceptors. In contrast, staining IMPG2[cK0]-expressing photoreceptors for IMPG2^m^ revealed a pronounced OS signal in addition to the IS signal. Measurement of the ratio of IMPG2^m^ in OS vs. IS revealed a significant accumulation of IMPG2^m^ in the OS when its ubiquitination is blocked (**Fig. 7C**).

**Figure 7:**
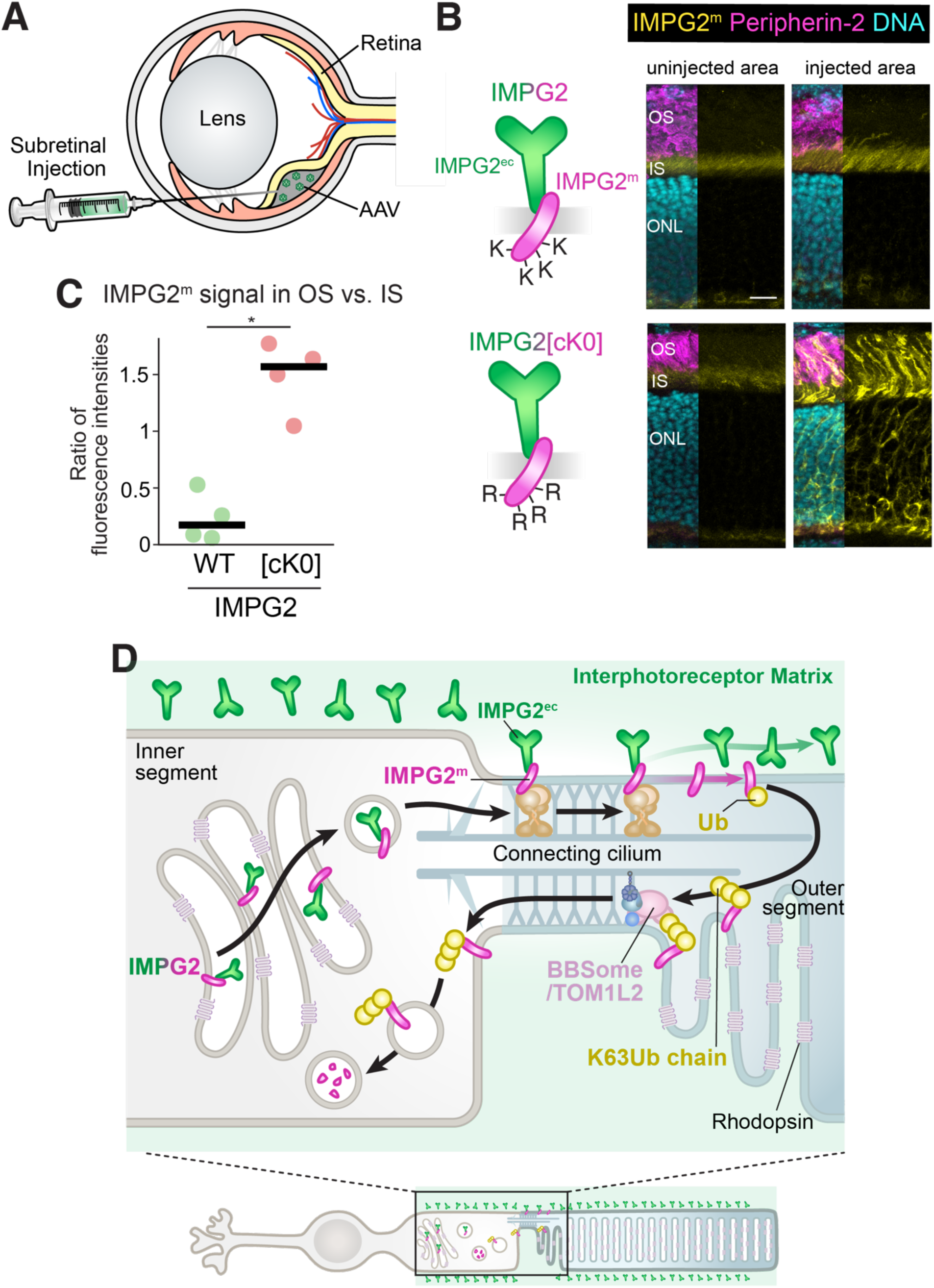
IMPG2 ubiquitination is required for the retrieval of IMPG2^m^ from outer segments. **A.** Schematic of the subretinal injection strategy used to deliver IMPG2 or a variant with all cytoplasmic lysines mutated to arginines (IMPG2[cK0]) into photoreceptors. Constructs expressing IMPG2 variants under control of the rhodopsin kinase promoter were packaged into pseudotyped AAV5 and injected into the subretinal space at P2. Retinal sections were analyzed three weeks after injection. **B-C.** Mutation of the cytoplasmic lysines in IMPG2 blocks exit of IMPG2^m^ from the OS. **B.** Representative images of uninjected and injected regions stained for IMPG2^m^ (yellow), peripherin (magenta), and DNA (cyan). See **Fig. S5** for additional examples from another injection. Scale bar: 10 μm. **C.** The fluorescence intensities for IMPG2^m^ were measured in the OS and IS in the injected area, and the ratios of OS-to-IS intensities plotted as dot plots. *N* = 1 retina each from 4 mice with 1 section per animal. Statistical significance was evaluated by an unpaired Mann-Whitney test (*, *p* < 0.05). **D.** Model of constitutive IMPG2 entry into the OS and subsequent BBSome- and ubiquitin-mediated retrieval of IMPG2^m^.

The similarity in the staining pattern of IMPG2^m^[cK0] in wild-type photoreceptors and IMPG2^m^ in *Bbs4^−/−^* photoreceptors suggests that IMPG2 ubiquitination and the BBSome are both required for the retrieval of IMPG2^m^ from the OS. A key difference between the staining pattern of IMPG2^m^[cK0] in wild-type photoreceptors and IMPG2^m^ in *Bbs4^−/−^* photoreceptors is the increased signal of IMPG2^m^[cK0] in the cell soma. This difference suggests that the ubiquitination of IMPG2^m^ is required not just for its retrieval from the OS but also for its degradation in the cell body, presumably via sorting to the lysosome.

The photoreceptor matrix consisting of IMPG1, IMPG2^ec^, and associated glycans is most enriched around the OS^46,51,52^. A simple model for delivery of IMPG2^ec^ to the milieu surrounding the OS holds that IMPG2 traffics to the OS where it undergoes cleavage and shedding of IMPG2^ec^, and IMPG2^m^ is then returned to the IS for degradation. We propose that after separating from IMPG2^ec^, IMPG2^m^ undergoes K63-liked ubiquitination which engages this important cargo onto the BBSome for removal from the OS (**Fig. 7D**).

## DISCUSSION

### Matrix deposition and membrane fragment turnover drive the physiological cycling of IMPG2 between inner and outer segment

Our findings uncover a constitutive membrane recycling pathway in photoreceptors, wherein the BBSome selectively retrieves the transmembrane fragment of IMPG2 (IMPG2^m^) from the OS through K63-linked ubiquitination. This mechanism departs from prevailing models that primarily frame BBSome function as a corrective system for mislocalized membrane proteins in photoreceptors, or as an on-demand device to retrieve GPCRs from conventional cilia in an activity-dependent manner. Instead, our data reveal that the BBSome is essential for normal photoreceptor homeostasis by mediating the physiological turnover of endogenous cargo.

IMPG2 is a photoreceptor-specific proteoglycan that delivers its large extracellular domain (IMPG2^ec^) to the interphotoreceptor matrix^45–47,51^. Because most of IMPG2^ec^ is found around the OS^46,51,52^, we propose that the full-length IMPG2 precursor is targeted to the OS, where autoproteolytic cleavage of IMPG2^46^ releases IMPG2^ec^ from the residual membrane-bound fragment IMPG2^m46^. Our data show that IMPG2^m^ is ubiquitinated in the OS and returned to the IS via a BBSome- and TOM1L2-dependent retrieval pathway. Denaturing affinity purification and K63 linkage-specific probes demonstrate that IMPG2^m^ accumulates in polyubiquitinated forms in *Bbs4*^−/−^ OSs, and disruption of either BBSome function or IMPG2 ubiquitination leads to the robust accumulation of IMPG2^m^ in the OS. These findings indicate that IMPG2^m^ does not passively leak in the OS; rather, it undergoes constitutive cycling as part of a tightly regulated renewal process for components of the photoreceptor matrix. The early onset and steady-state accumulation of K63-linked ubiquitin in OSs lacking BBSome function, observable as early as P8 (**Fig. 2**), further supports the idea that BBSome-mediated cargo retrieval is intimately associated with physiological protein turnover and not merely correcting accidental leakage into the OS.

The increased signal of IMPG2^m^[cK0] not only in the OS but also in the IS and the ONL (**Fig. 7B**) indicates that ubiquitination of IMPG2^m^ is required for both its retrieval from the OS and its degradation within the photoreceptor soma. Taken together, our data position K63-linked ubiquitination as a decisive signal for BBSome-mediated retrieval and overall clearance in photoreceptors (**Fig. 7D**).

### The role of the BBSome in photoreceptors: quality control of the OS proteome or constitutive cycling?

The SNARE complex composed of syntaxin-3 and its partners Munc18-1 (syntaxin-binding protein 1, STXBP1) and synaptosomal-associated protein 25 (SNAP25), as well as several other proteins, accumulates in the OS of transition zone mutants^20,21^. Because these proteins also accumulate in OSs when BBSome function is disrupted^15,40^, it has been proposed that their normally slow leakage into the OS is corrected by the BBSome^17,22^. One puzzling aspect of this model is the requirement for these unrelated proteins to carry some feature marking them as non-OS to enable their selection by the retrieval machinery. An additional unanswered question is that a SNARE complex with synaptic functions would be particularly prone to leakage when most IS proteins remain well-retained.

An alternative interpretation considers that the BBSome physically engages transition zone components when ferrying cargoes through the transition zone^53,54^. First, single-molecule imaging of BBSome cargoes revealed a strict dependence on the BBSome for their transition zone crossing^55^. Second, the BBSome physically interacts with transition zone proteins^56^. Third, multiple genetic interactions between the BBSome and transition zone proteins have been reported^57–60^. Together, these findings suggest that transition zone mutants may be defective in assisting BBSome crossing rather than by creating a leakier sieve.

In this context, we propose that constitutive cycling between IS and OS underlies the previously observed accumulations of syntaxin-3 –and other proteins– in BBSome-defective OSs. Given its role in mediating the fusion of rhodopsin carrier vesicles with the periciliary ridge complex at the base of the connecting cilium^38,39^, syntaxin-3 and its SNARE partners may enter the cilium after vesicle fusion. Alternatively, syntaxin-3 associates with the OS disk rim proteins peripherin 2 and ROM1 (retinal outer segment membrane protein 1)^61^, and may function either in the targeting of these tetraspanins to the OS^61,62^ or by transiently assisting their function in disk morphogenesis. In support of syntaxin 3 undergoing constitutive entry into the OS, discrete signals of syntaxin-3 have been detected at the base of the mouse OS^61^ and most of syntaxin-3 localizes to the OS in human photoreceptors^63^. Furthermore, syntaxin-3 already accumulates markedly in the OS of *Bbs8^−/−^* mouse photoreceptor at P10, a very early stage of OSs biogenesis^26^. Thus, rather than accidently leaking into the OS, syntaxin-3 may traffic to the OS as part of its functional repertoire.

Finally, although syntaxin-3 was enriched in *Bbs4*^−/−^ OSs and identified in the UbK63-associated proteome, it did not show increases in covalent ubiquitination and did not respond to manipulations of ciliary UbK63 levels in IMCD3 cells. These observations indicate that syntaxin-3 accumulates in cilia and OSs of BBSome-defective cells independently of its ubiquitination status and may not represent a direct cargo of the BBSome. One possibility is that the previously observed alterations in lipid composition in the OS of Bbs mutants^29^ indirectly trap syntaxin-3 in OS.

### Molecular basis of photoreceptor degeneration in Bardet-Biedl Syndrome

The BBSome cargoes and constitutive cycling mechanism that we uncovered have direct implications for understanding the retinal pathophysiology of BBS. First, failed clearance of specific membrane proteins from the OS may directly poison photoreceptors. Given that IMPG2^ec^ is abundant in the photoreceptor matrix, the excessive amounts of IMPG2^m^ in the membrane of *Bbs* OSs may adversely affect OS membrane properties. Secondly, defective retrieval of IMPG2^m^ may compromise homeostasis of the interphotoreceptor matrix. Because mutations in IMPG2 cause retinal degeneration^64^ and in light of the broad functional importance of the interphotoreceptor matrix in retinal physiology^45^, even subtle defects in IMPG2^ec^ deposition defects may trigger retinal degeneration in BBS. Finally, the buildup of K63-linked ubiquitin chains in OS membranes may deplete free ubiquitin pools and lead to secondary defects in proteostasis, as previously noted in *Bbs12*-deficient photoreceptors^65^.

In summary, our study shows that the BBSome is integral to the constitutive turnover of membrane proteins in photoreceptors. Rather than merely acting as a quality control system to clear mislocalized proteins from the OS, the BBSome supports photoreceptor homeostasis by retrieving physiological cargoes marked by K63Ub chains.

## ACKNOWLEDGEMENTS

We thank Mark Hochstrasser, Rohan Baker, David Komander, for the gift of plasmids; Val Sheffield for the gift of *Bbs4* mouse strains; Suling Wang for help with graphic design; Robyn Eisert with technical assistance with the TMT analysis, Arlene Drack, Seongjin Seo and Val Sheffield for advice on subretinal injection and Steven Fliesler and Ezequiel Salido and all members of the Nachury lab for stimulating discussions. This work was funded by NIH (GM089933 and EY031462 to MVN). This work was made possible, in part, by EY002162 - Core Grant for Vision Research and by the Research to Prevent Blindness Unrestricted Grant (MVN). MK is a CPRIT scholar in Cancer Research (RR220032).

---

**Supplementary Figure 1:**
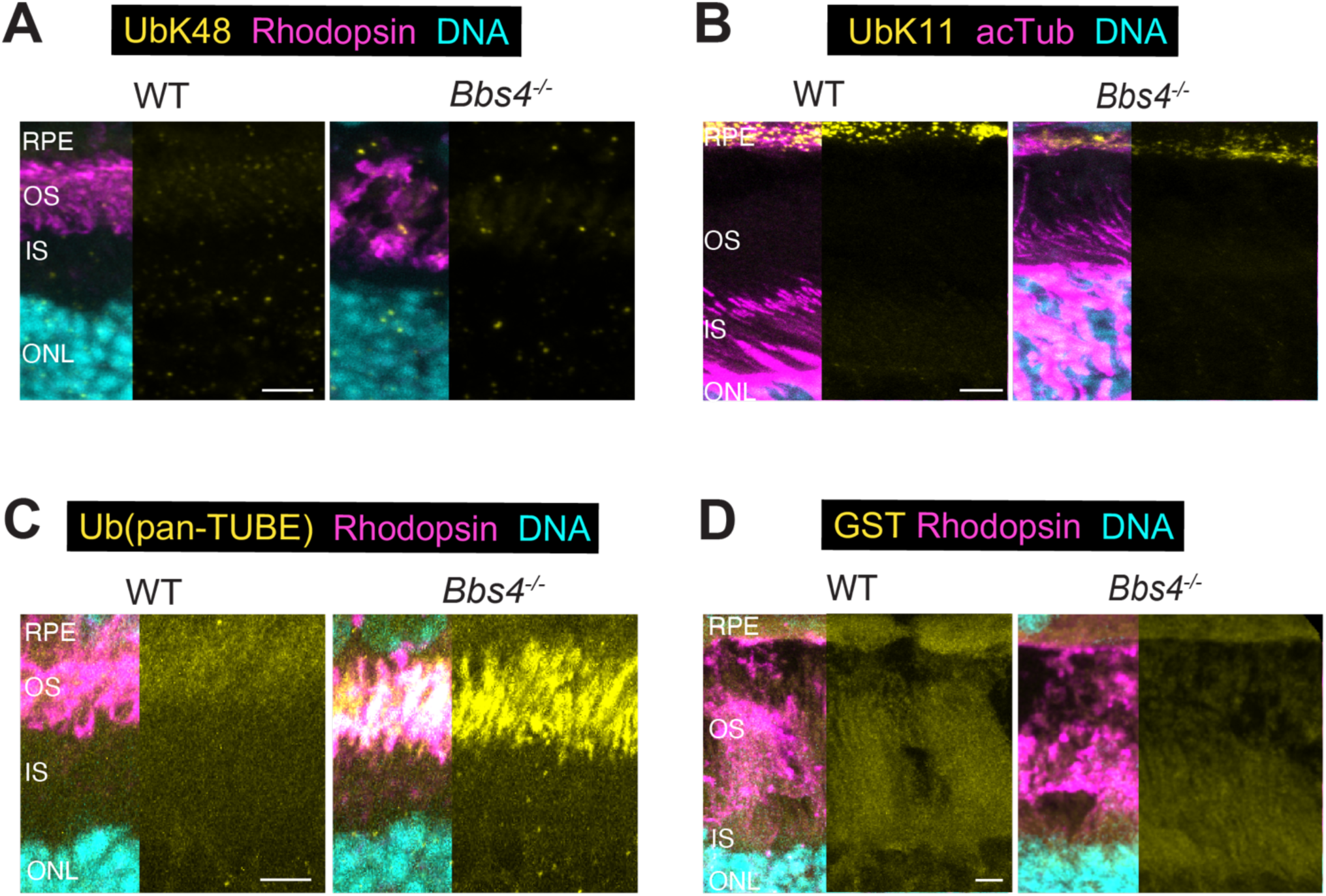
Localization of ubiquitin chains in wild-type and *Bbs4^−/−^* photoreceptors. Related to Figures 1 and 2. **A-D.** Representative images of retinal cryosections from wild-type and *Bbs4^−/−^* mice stained for various ubiquitin chain linkages and controls. (**A**) K48-linked ubiquitin chains (UbK48, mAb Apu2, yellow), rhodopsin (magenta), and DNA (cyan) in P15 animals. (**B**) K11-linked ubiquitin chains (UbK11, mAb2A3/2A6, yellow), acetylated tubulin (acTub, magenta) and DNA (cyan) in P18 animals. (**C**) all Ub linkages (GST-panTUBE, yellow), rhodopsin (magenta), and DNA (cyan) in P15 animals. (**D**) GST-only control (yellow), rhodopsin (magenta), and DNA (cyan) in P15 animals. OS: outer segment, IS: inner segment, ONL: outer nuclear layer, RPE: retinal pigmented epithelium. Scale bars: 5 μm.

**Supplementary Figure 2:**
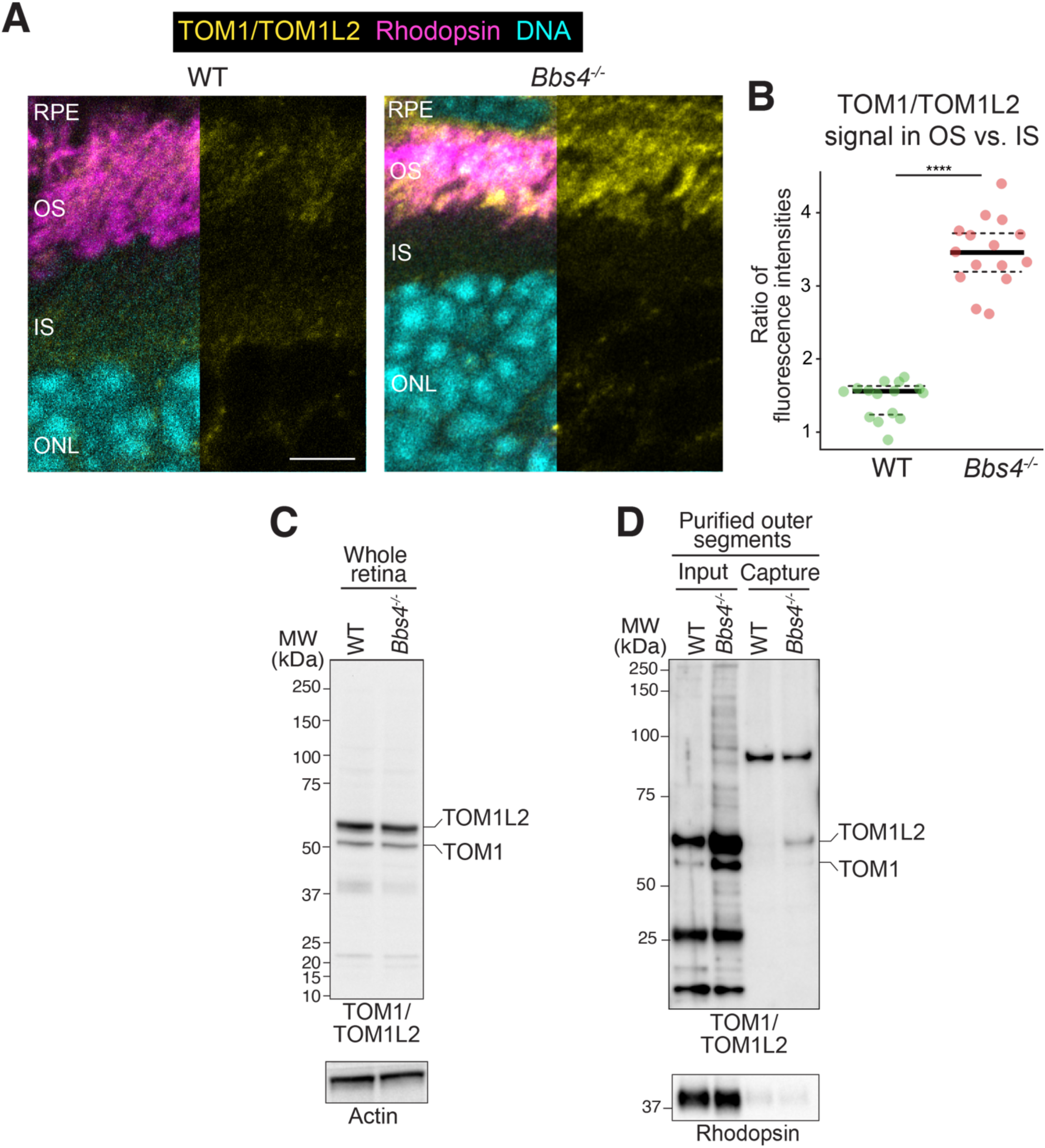
TOM1/TOM1L2 accumulation in the OS of *Bbs4^−/−^* photoreceptors. Related to Figure 2. **A**. Representative images of retinal cryosections from wild-type and *Bbs4^−/−^* mice at P15 stained for TOM1/TOM1L2 (yellow), rhodopsin (magenta), and DNA (cyan). Scale bar: 5 μm. **B**. Quantitation of TOM1/TOM1L2 fluorescence intensity in outer and inner segments at P15. Ratios of outer to inner segment fluorescence intensities are plotted as dot plots (*N* = 2 retinas each from 3 mice with 3 retinal cross-sections per animal). Statistical significance was determined by an unpaired non-parametric Mann-Whitney test (****, *p* ≤ 0.0001). **C**. Immunoblot of whole retinal lysates from wild-type and *Bbs4^−/−^* mice at P15 probed for TOM1/TOM1L2 (top panel) and actin (loading control, bottom panel). The commercial antibody (Abcam ab96320) raised against TOM1 detects both TOM1 and TOM1L2^14,66^. **D**. GST-K63TUBE eluates from wild-type and *Bbs4^−/−^* OSs captures were probed for TOM1/TOM1L2 (top panel) and rhodopsin (bottom panel). 0.1% of inputs and 20% of eluates were loaded for immunoblotting.

**Supplementary Figure 3:**
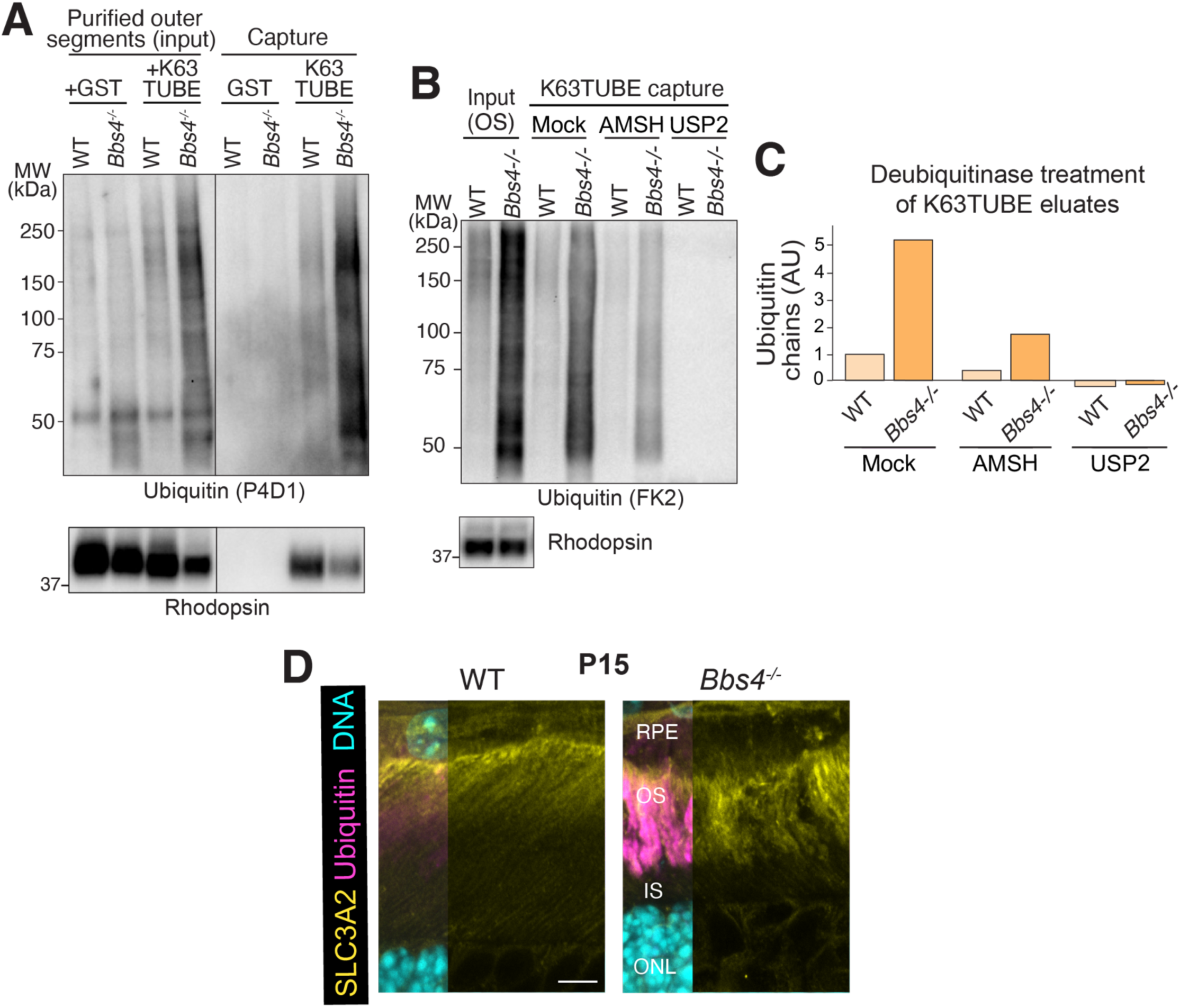
K63TUBE captures, deubiquitinase assays and SLC3A2 staining. Related to Figures 3 and 4. **A**. K63TUBE protects UbK63 chains and captures K63Ub chains from photoreceptor OSs. Purified OSs from P15 wild-type and *Bbs4^−/−^* mouse retinas were mixed with GST or GST-K63TUBE, lysed and captured as illustrated in Fig. 4A. Samples were probed for ubiquitin (P4D1 antibody, top panel) and for rhodopsin (loading control, bottom panel). 0.1% of inputs and 20% of eluates were loaded for immunoblotting. **B-C.** Deubiquitinase-based analysis of ubiquitin chain architecture in the K63TUBE eluates. **B.** K63TUBE eluates from wild-type or *Bbs4^−/−^* OS captures treated with AMSH (K62 linkage-specific) or USP2 (all linkages) were probed for ubiquitin (FK2). **C.** Quantitation of integrated intensity above 37 kDa from (**B**), plotted in a bar graph. **D.** Accumulation of SLC3A2 in *Bbs4^−/−^* outer segments. Representative images of P15 retinal sections stained for SLC3A2 (yellow), ubiquitin (magenta) and DNA (cyan). Scale bar: 5 μm. RPE: retinal pigmented epithelium, OS: outer segment, IS: inner segment, ONL: outer nuclear layer.

**Supplementary Figure 4:**
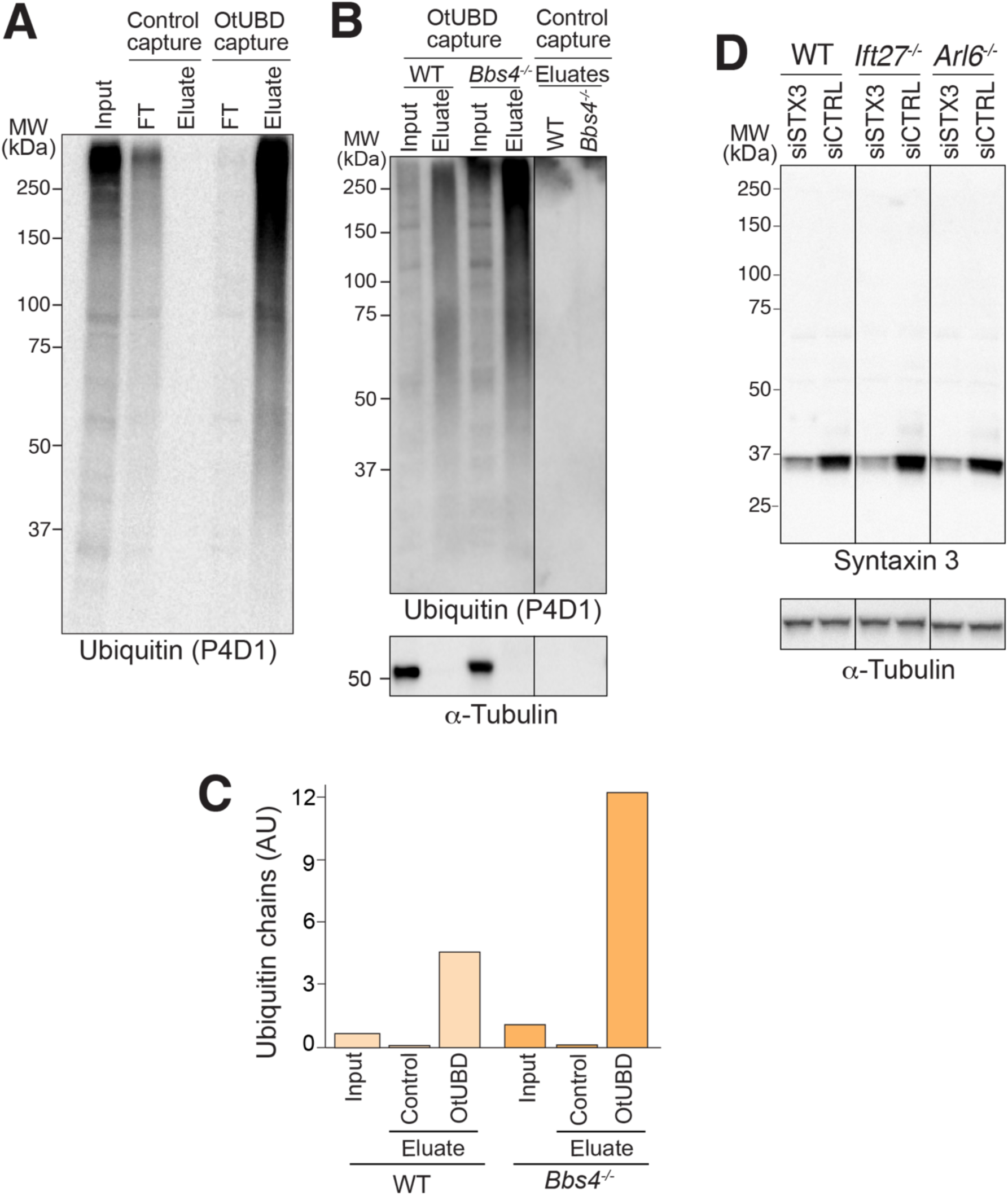
Validations of OtUBD capture strategy and syntaxin-3 antibody. Related to Figures 5 and 6. **A.** OtUBD captures ubiquitinated proteins from HEK293T cells. Cells were treated with MG-132 for 4 hr. Lysates were prepared under denaturing conditions and captured on OtUBD resin or control resin as illustrated in Fig. 5A. 4% input and flowthrough (FT) and 15% eluate fractions were probed for ubiquitin (P4D1). **B-C.** OtUBD captures ubiquitinated proteins from photoreceptor OSs. **B.** Extended blot from Fig. 5B. Purified OSs (input), OtUBD and control eluates were probed for ubiquitin (P4D1, top panel) or tubulin (loading control, bottom panel). 2% of inputs and 15% of eluates were loaded for immunoblotting. **C.** Quantitation of integrated intensity above 37 kDa from (**B**), plotted in a bar graph. **D.** WT, *Ift27^−/−^* and *Arl6^−/−^* IMCD3 cells were transfected with siRNAs targeting STX3 (siSTX3) or control siRNA (siCTRL). Lysates were probed for syntaxin-3 (top panel) or tubulin (loading control, bottom panel). Depletion efficiency was 54% in WT cells, 66% in *Ift27^−/−^* cells and 63% in *Arl6^−/−^* cells.

**Supplementary Figure 5:**
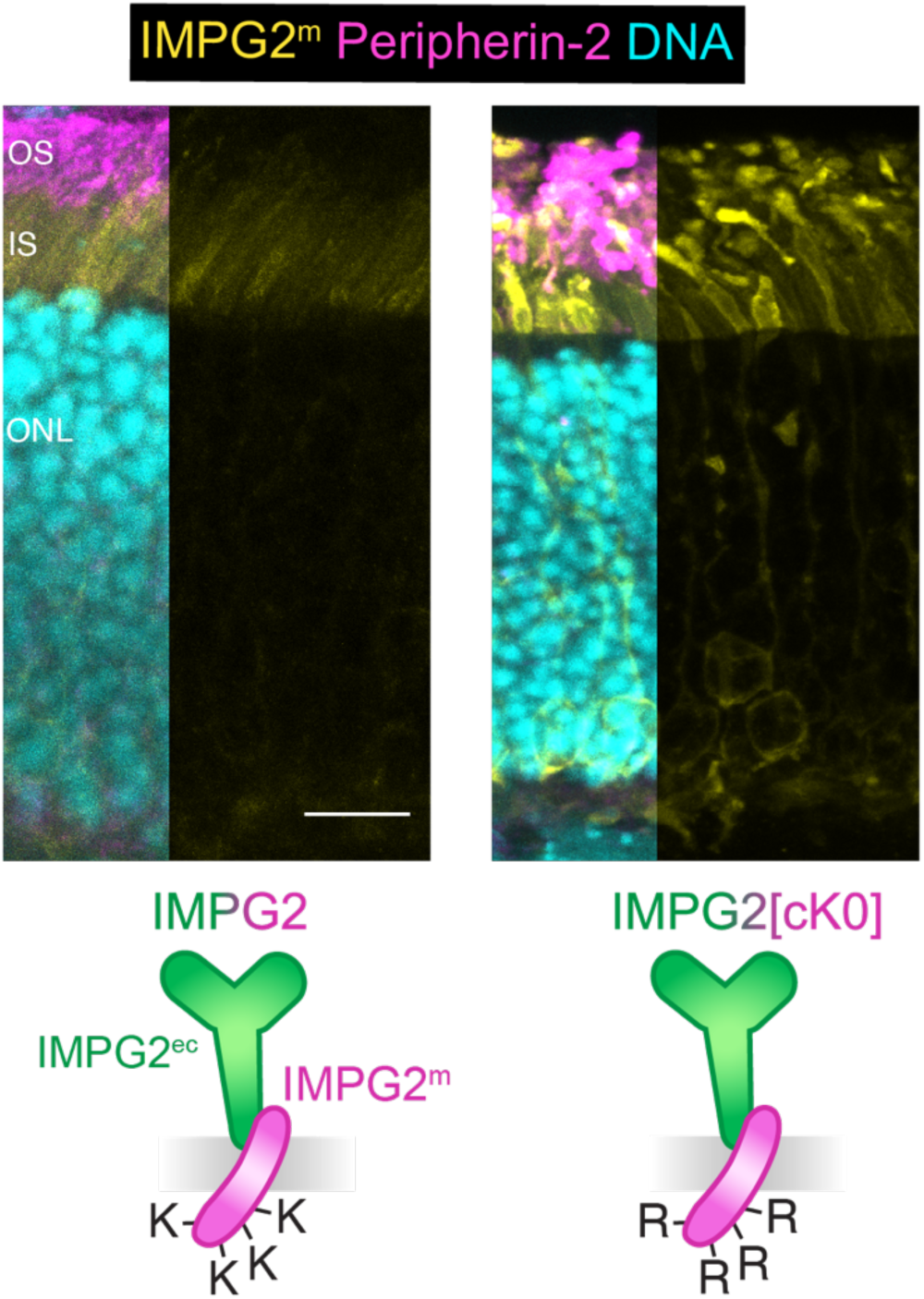
IMPG2^m^ ubiquitination is required for its retrieval from outer segments. Related to Figure 7. Expression constructs for IMPG2 variants packaged into AAV5 viruses were injected into the subretinal space as illustrated in Fig. 7A at P2. Representative images of injected regions from another animal than shown in Fig. 7B, stained for IMPG2^m^ (yellow), peripherin-2 (magenta), and DNA (cyan). OS: outer segment, IS: inner segment, ONL: outer nuclear layer. Scale bar: 5 μm.

## METHODS

Reagents are listed in Table S1

## DATA AND CODE AVAILABILITY

The mass spectrometry proteomics data have been deposited to the ProteomeXchange Consortium (http://proteomecentral.proteomexchange.org) via the PRIDE^67^ partner repository with the dataset identifier PXD065467. Other data reported in this paper will be shared by the lead contact upon request. This paper does not report original code.

## EXPERIMENTAL MODEL AND SUBJECT DETAILS

All IMCD3 cell lines used in the study were derived from a parental IMCD3-FlpIn cell line described previously^68^. IMCD3 cells were cultured in DMEM/F12 supplemented with 10% FBS at 37 °C with 5% CO_2_. Ciliation was induced by serum starvation of cells in serum-free medium for 16 to 24 hr. All cell lines were confirmed to be mycoplasma-free by monthly testing with the MycoAlert system (Lonza).

All mouse work adhered to the Guide for the Care and Use of Laboratory Animals from NIH and the ARVO Statement for the Use of Animals in Ophthalmic and Vision Research. Procedures were reviewed and approved by the University of California, San Francisco Institutional Animal Care and Use Committee (IACUC). Animals were housed in an AAALAC-accredited, specific-pathogen-free barrier facility under a 12 hr light / 12 hr dark cycle with irradiated chow and reverse-osmosis water provided ad libitum. Mice were examined daily, weighed weekly, and euthanized by carbon-dioxide inhalation followed by cervical dislocation when predefined humane end-points were reached; all efforts were made to minimize pain and distress. Both male and female mice were included in every experiment, and no sex-dependent differences were observed in any phenotype reported. The *Bbs4^−^* line was a gift from the Sheffield laboratory and maintained on a CD-1 background. *Bbs4* mice were genotyped as described^25^ (JAX protocol # 23573). Experimental *Bbs4^−/−^* and wildtype littermate control animals were generated by crossing heterozygous *Bbs4^+/−^* breeders. Outbred CD-1 mice (Charles River Laboratories, strain code 022) were obtained commercially and maintained under recommended husbandry conditions.

Zebrafish (Danio rerio) were maintained as described^69^. Embryos were raised at 28 °C in embryos medium. The *bbs1* line was previously described^29^. Zebrafish husbandry and experimental procedures were performed in accordance with Swiss animal protection regulations (Veterinäramt Zürich, Tierhaltungsbewilligung 150 and Tierversuch ZH116/2021-33632).

## METHOD DETAILS

### Plasmids

The coding sequences for the previously described K63TUBE (A7(Vx3)^70^) and pan-TUBE (hRAD23A UBA^71^) were gene-synthesized and cloned into pGEX6P vectors by Genscript.

The adenovirus-associated virus (AAV) shuttle plasmids for expression of full-length mouse IMPG2 (NM_174876.3) or IMPG2[cK0] (K1128R, K1160R, K1177R, K1212R) fused at the C-terminus to FLAG and T2A epitopes under the control of the rhodopsin kinase promoter (pAAV5-pRhoK-mIMPG2-FLAG-T2A and pAAV5-pRhoK-mIMPG2[cK0]-FLAG-T2A) were cloned by Vector Builder using gene synthesis, site-directed mutagenesis and conventional cloning.

### Antibodies

The following monoclonal antibodies were used for immunofluorescence: anti-rhodopsin (1D4, 1:1000), anti-acetylated tubulin (6-11-B, 1:500), anti-ubiquitin (FK2, 1:100), anti-lysine 48 ubiquitin linkage (Apu2, 1:100), anti-lysine 11 ubiquitin linkage (2A3/2A6, 1:50), anti-JAMB (150005, 1:100), anti-SLC3A2 (RL388, 1:100), anti-peripherin-2 (2B7, 1:500, **Figs.7B and S5**), anti-FLAG (M2, 1:500).

The following polyclonal antibodies were used for immunofluorescence: anti-peripherin-2 (1:500), anti-IMPG2^m^ (1:100), anti-syntaxin3 (1:100), anti-TOM1/TOM1L2 (1:500).

The following secondary antibodies were used for immunofluorescence: Alexa Fluor 488-, 568-, and 647-conjugated anti-rabbit IgG (Invitrogen, 1:500), Alexa Fluor 488-, 568-, and 647-conjugated anti-mouse IgG (Invitrogen, 1:500), Alexa Fluor 647-conjugated anti-rat (Invitrogen; 1:100), Cy3-conjugated anti-mouse IgG2b, Alexa Fluor 488-conjugated anti-mouse IgG1, Cy5-conjugated anti-rabbit IgG (Jackson ImmunoResearch, 1:500).

The following antibodies were used for immunoblotting: anti-ubiquitin (P4D1, 1:1000, **Figs. 5, S3A, S4**), anti-ubiquitin (FK2, 1:1000, **Figs. 1, 3, S3B**) anti-rhodopsin (1D4, 1:1000), anti-lysine 63 ubiquitin linkage (Apu3, 1:1000), anti-IMPG2^m^ (1:1000), anti-syntaxin3 (1:1000), anti-TOM1/TOM1L2 (1:1000), anti-α-tubulin (1:2000), anti-actin (1:1000).

### Cell Culture Methods

#### Cell culture

A parental IMCD3-FlpIn cell line (gift from Peter K. Jackson, Stanford University, Stanford, CA) was modified to generate all stable cell lines used in the study. IMCD3-FlpIn cells were cultured in DMEM/F12 (11330-057; Gibco) supplemented with 10% FBS (100-106; Gemini Bio-products), 100 U/mL penicillin-streptomycin (400-109; Gemini Bio-products), and 2 mM L-glutamine (400-106; Gemini Bio-products). The IMCD3 *Ift27^−/−^* and *Arl6^−/−^* cell lines were described previously^72^. Ciliation was induced by serum starvation in media containing 0.2% FBS for 16 to 24 hr.

Human epithelial kidney (HEK293) cells were cultured in high-glucose DMEM (11965-092; Gibco) supplemented with 10% FBS, 5% MEM-NEAA (11140-050; Gibco), 5% sodium pyruvate (11360-070; Gibco) in addition to penicillin-streptomycin.

#### Transfection of cultured cells

For siRNA treatments, 50,000 WT, *Arl6^−/−^* or *Ift27^−/−^* IMCD3 cells were transfected with siRNA duplexes targeting mouse syntaxin-3 or control siRNAs using Lipofectamine RNAiMAX (13-778-030, Thermo Scientific) via reverse transfection in a 24-well plate following the manufacturer protocol. In brief, 3µL of RNAiMAX were diluted in 100 µl of Opti-MEM (31985070, Life Technologies) and incubated at room temperature for 5 min. Next, 1 µl of siRNA solution (10µM stock) was added to the diluted transfection reagent and incubated for 20 min before mixing with the cells in suspension. Cells were serum-starved for 48 hr after transfection and cultured for another 16 hr. Cells were lysed with RIPB buffer and equivalent amounts of clarified supernatants immunoblotted for syntaxin-3 or α-tubulin.

#### Imaging of cultured cells

IMCD3 cells were fixed with 4% paraformaldehyde (50-980-487, Thermo Fisher Scientific) in PBS for 15 min at 37 °C. Cells were permeabilized in IF buffer [PBS supplemented with 0.1% Triton X-100, 5% normal donkey serum (017-000-121; Jackson ImmunoResearch Laboratories), and 3% bovine serum albumin (BP1605-100; Thermo Fisher Scientific)]. Cells were then incubated at room temperature for 1 hr with primary antibodies diluted in IF buffer, washed three times in IF buffer, and incubated with secondary antibodies (Alexa Fluor-conjugated fluorophores) diluted in IF buffer for 30 min. Cells were washed three times with IF buffer and DNA stained with Hoechst 33258 (H1398; Molecular Probes). Cells were washed twice with PBS, and coverslips mounted on slides using Fluoromount-G (17984-25; Electron Microscopy Sciences). Cells were imaged on a LSM 900 confocal microscope (Zeiss) equipped with a Plan-Apochromat 63X/1.40 Oil DIC M27oil objective. Z-stacks were acquired at 0.2 μm interval.

#### Image analysis of cultured cells

Maximum intensity projections were generated from the consecutive Z-sections where cilia were in focus in the acetylated tubulin channel. Next, masks were generated for each cilium by drawing a 3-pixel wide segmented line along each cilium in the acetylated tubulin channel. The masks were transferred to the syntaxin-3 channel and background-corrected ciliary fluorescence intensities (F_cilia_) were calculated by subtracting from the raw integrated density of cilia fluorescence (F_raw_) the integrated density of an adjacent area (F_background_). Ciliary intensities (F_cilia_= F_raw_ - F_background_) of at least 50 cilia per condition were collected and plotted as dot plots using PlotsofData^73^. Statistical analyses were performed as indicated in the respective figure legends using GraphPad Prism on the combined dataset from at least 3 experiments^73^.

### Biochemical Methods

#### Protein expression and purification

GST-tagged TUBE proteins were expressed from pGEX6P vectors in Rosetta2(DE3)-pLysS cells grown in 2xYT medium (Millipore Sigma, Y2627) at 37 °C until the optical density (OD) at 600 nm reached 0.6. Protein expression was then induced with 0.2 mM IPTG at 18 °C for 16 hr. After induction the cells were pelleted down at 6,000 x *g* for 15 min at 4 °C and pellets were resuspended in 2XT buffer (40 mM Tris pH 7.4, 200 mM NaCl, 5 mM DTT) supplemented with protease inhibitors (10 μg/mL each of Leupeptin, Pepstatin A, Bestatin, 0.8 mM Aprotinin, 1 mM AEBSF and 15 mM E-64) and lysed by sonication. The lysates were clarified by centrifugation at 30,000 x *g* for 30 min at 4 °C. The clarified lysates were loaded onto Glutathione Sepharose 4B (Cytiva) and the recombinant proteins eluted with 50 mM Glutathione, pH 9.0 in 2XT buffer. The eluted proteins were then dialyzed against XT buffer (20 mM Tris pH 7.4, 100 mM NaCl, 1 mM DTT) with one change of buffer and flash-frozen in liquid nitrogen after addition of 5% (w/v) glycerol.

His_6_-AMSH and His_6_-USP2cc were expressed from pOPINB-AMSH* and pET15b-USP2cc following the same protocol as the GST-TUBEs. Bacterial pellets were resuspended in lysis buffer (20mM Tris pH 8.9, 500mM NaCl, 50mM imidazole, 0.5mM DTT) supplemented with protease inhibitors (10 μg/ml each of Leupeptin, Pepstatin A, Bestatin, 0.8mM Aprotinin, 1mM AEBSF and 15mM E-64) and lysed by sonication. His_6_-tagged proteins were purified on Ni^2+^-NTA resin (Qiagen;). Elutions were performed in lysis buffer supplemented with 400 mM imidazole. For AMSH, the His_6_-tag was cleaved with GST-HRV3C. Following the removal of GST-HRV3C from eluate fraction via capture on Glutathione Sepharose 4B resin (Cytiva), AMSH was further purified on Superdex 75 (Cytiva) in 20 mM Tris pH 8.0, 150 mM NaCl, 5 mM ß-mercaptoethanol. Purified recombinant proteins were flash-frozen in liquid nitrogen after addition of 5% (w/v) glycerol.

OtUBD recombinant protein was expressed from pET21a-cys-His6-OtUBD as described^49^. The OtUBD resin and the negative control resins were then prepared by conjugating OtUBD or free cysteine on SulfoLink resin (Thermo, 20401).

#### Enrichment of mouse photoreceptor outer segments

Mice were killed by CO_2_ asphyxiation followed by cervical dislocation. The skin around the eye socket was held so the eyeball protruded. A horizontal cut was made across the cornea using a razor blade. Curved forceps were inserted under the eyeball, gently squeezing to remove the lens. Continued squeezing and pulling with the forceps is used to remove the vitreous humor. Further squeezing and pulling detaches the retina from the eye cup. Each retina was directly transferred into a tube containing buffer. Retinas were gently dounced with a micro-pestle (USA Scientific 14155290) in a 1.5 mL microfuge tube, and large debris were spun down at 100 x *g* for 3 minutes at 4 °C. The outer segment fraction was lysed in TUBE lysis buffer (see below) overnight at 4 °C. Lysates were clarified by centrifugation at 12,000 x *g* for 20 minutes at 4 °C and the clarified supernatants used in downstream analyses.

#### TUBE captures

Enriched photoreceptor outer segments from WT and *Bbs4^−/−^* mice were lysed in TUBE lysis buffer (20 mM sodium phosphate pH 6.8, 50 mM NaF, 5 mM NaPPi, 1% NP-40, 2 mM EDTA, protease inhibitors (10 μg/mL each of Leupeptin, Pepstatin A, Bestatin, 0.8 mM Aprotinin, 1 mM AEBSF and 15 mM E-64)) supplemented with 5.8µM GST-TUBE or GST alone as described^71^. The outer segment/GST-TUBE mixtures were then purified over Glutathione Sepharose 4B resin. Following three washes with PBST (PBS, 0.05% Tween-20), bound proteins were eluted by cleavage with 1.5 µM GST-HRV3C in 100 µL PBST (1xPBS, 0.05% Tween-20) overnight at 4 °C. The cleaved eluates were analyzed by western blotting or mass spectrometry.

#### Whole retinal extract preparation

Six mouse retinas from WT and *Bbs4^−/−^* mice freshly dissected as above were directly lysed in 100 µL RIPA buffer (20 mM Tris-HCl pH 7.5, 150 mM NaCl, 1 mM EGTA, 1 mM EDTA, 1%NP-40, 1% sodium deoxycholate, 2.5 mM NaPPi, 1 mM β-glycerophosphate, 1 mM Na_3_VO_4_) supplemented with protease inhibitors (10 μg/mL each of Leupeptin, Pepstatin A, Bestatin, 0.8 mM Aprotinin, 1 mM AEBSF and 15 mM E-64). Retinas were manually lysed with a microfuge pestle (USA Scientific 14155290), and incubated on ice for 30 minutes. The lysates were then vortexed briefly and clarified by centrifugation at 12, 000 x *g* for 10 minutes at 4 °C. Equivalent amounts of the clarified supernatant for each genotype were immunoblotted for TOM1/TOM1L2 and actin.

#### Immunoblotting and analyses

Protein samples were resolved on 4-12% NuPAGE (Thermo Fisher) or SurePAGE (Genscript) Bis-Tris gels and transferred onto polyvinylidene fluoride membrane blots. Membranes were probed overnight at 4 °C with primary antibodies diluted in blocking buffer (10% Fish Blocking Serum (ThermoFisher; 37527) in TBST), and incubated with HRP-conjugated secondary antibodies (Jackson Immunoresearch) in TBST for 1 hr at room temp. Signals were developed with ECL Substrate (Thermo Scientific, 32106), and chemiluminiscent signals imaged on a ChemiDoc imaging system (Bio-Rad). Band and smear intensities were quantified in ImageLab v6.1 (Bio-Rad). All densitometry values were background-corrected against an empty lane and normalized to the appropriate loading control bands. For ubiquitin smear densitometric analysis, the integrated density of each lane from 37 kDa to the top of the blot was measured after background subtraction of an equally sized region from an adjacent empty lane. Ubiquitinated IMPG2 smears (**Fig.5 C,D**) were quantified by measuring the integrated density signal from 37 kDa (above the 32 kDa IMPG2^m^ band) to the top of the membrane.

#### *In vitro* deubiquitination assay

*In vitro* deubiquitination reactions were carried out on eluates from K63TUBE purifications of WT and *Bbs4^−/−^* outer segment lysates as described^33^. Briefly, recombinant AMSH or USP2cc were diluted to a final concentration of 2.5 µM in 25 mM Tris-HCl pH 7.5, 150 mM NaCl, and 10 mM DTT and preincubated at 23 °C for 10 min. 24 µL K63TUBE eluates were mixed with 3 µL of diluted DUB (250 nM final) and 3 µL 10X DUB Buffer (500 mM Tris-HCl pH 7.5, 500 mM NaCl, and 50 mM DTT) and reactions conducted at 37 °C for 2 h. Reactions were terminated by the addition of 4× LDS sample buffer and immediate heating to 95 °C for 5 min.

#### Capture of ubiquitinated proteins under denaturing conditions

Isolation of ubiquitinated proteins with OtUBD resin was performed under denaturing conditions as described^49^. Briefly, outer segment fractions were resuspended in lysis buffer (10 mM Hepes pH 7.4, 130 mM NaCl, 3.6 mM KCl, 2.5 mM MgCl₂, 1.2 mM CaCl₂, 0.02 mM EDTA, 1%NP-40, 5 mM N-ethylmaleimide, 1 mM PMSF) and 480 mg of urea was added for each mL and the resulting mixture was briefly vortexed to dissolve the urea and bring its concentration to 8 M final. Following incubation at room temperature for 30 mins, the lysates were clarified by centrifugation at 12,000 x g for 20 mins. Clarified lysates were diluted 1:1 with column buffer (50 mM Tris HCl pH7.5, 150 mM NaCl, 1 mM EDTA, 0.5 % Triton X-100, 10 % glycerol) to bring urea concentration to 4 M final. Equal protein amounts from wild-type and *Bbs4^−/−^* samples were incubated with OtUBD or control resins pre-equilibrated in column buffer at 4 °C for 2 hr. Resins were washed sequentially with 15 bed volumes each of column buffer containing 4 M urea, Wash Buffer I (50 mM Tris-HCl, 150 mM NaCl, 0.05 % Tween-20, pH 7.5), and Wash Buffer II (50 mM Tris–HCl, 1 M NaCl, pH 7.5). Bound proteins were eluted with 1x LDS sample buffer at 95 °C for 5 min.

For HEK-293T cell assays, confluent cultures were treated with 1 µM MG-132 for 4 hr prior to lysis in urea buffer. OtUBD and control captures were then performed as above.

### TMS-MS of TUBE captures

K63TUBE captures were conducted from WT and *Bbs4^−/−^* outer segment fractions with eight retinas per genotype in biological triplicates.

#### SP3 Bead Binding and On-Bead Digestion

GST and GST-TUBE eluates were processed by a modified SP3 protocol^74^. Each lysate was adjusted to 250 µL with 50 mM HEPES, 50 mM NaCl. DTT (1 M in water, freshly prepared) was added to 5 mM, and samples were shaken at 1,000 rpm for 30 min at 60 °C. After cooling to room temperature (RT), reduced proteins were alkylated in the dark for 30 min with 400 mM iodoacetamide in 50 mM ammonium bicarbonate (final 20 mM). Alkylation was quenched by adding DTT to 50 mM and incubating 15 min at RT. SP3 beads were prepared at 50 µg/µL in HPLC-grade water (1:1 hydrophobic/hydrophilic Sera-Mag SpeedBeads). 10 µL of the bead suspension were added to each sample, followed by 250 µL ethanol. After gentle vortexing, binding proceeded for 5 min at 24 °C, 1,000 rpm. Beads were collected on a magnetic rack and washed three times with 250 µL 80 % ethanol. Proteins were digested on-bead in 50 µL digestion buffer (200 mM EPPS, pH 8.5, 2 % acetonitrile, v/v) with Lys-C (2 mg/mL) at a 1:50 enzyme:substrate ratio for 3 hr at 37 °C. A further 50 µL digestion buffer and trypsin (2 mg/mL) were then added at a 1:100 ratio, and digestion continued 7 hr at 37 °C. Beads were removed magnetically and clear supernatants transferred to fresh tubes.

#### TMT Labeling

Digested peptides were brought to 30% (v/v) acetonitrile and labeled with TMTpro 16-plex reagent. Labeling was carried out at RT with brief vortexing every 10 min for 1 h. A pooled 12-plex aliquot analyzed by MS³ confirmed >95% labeling efficiency. Reactions were quenched with 50% hydroxylamine (final 0.5%, v/v) for 15 min, acidified to pH < 3 with formic acid, and combined into a 12-plex set. Solvent was evaporated under reduced pressure, and peptides fractionated with the Pierce High-pH Reversed-Phase kit into 12 fractions eluted at 10, 12.5, 15, 17.5, 20, 25, 30, 35, 40, 50, 65 and 80% acetonitrile. Fractions were merged (1 + 7, 2 + 8, 3 + 9, 4 + 10, 5 + 11, 6 + 12), dried, and cleaned by StageTip.

#### Mass Spectrometry

Peptides were separated on an EASY-nLC 1200 (Thermo Fisher) using a 30–40 cm, 100 µm-ID column packed with 2.6 µm Accucore resin and heated to 60 °C. At 450 nL/min, a 5–100% Buffer B gradient (240 min) was applied; Buffer A contained 0.125% formic acid (FA) and Buffer B 95% acetonitrile, 0.125% FA. Eluates were analyzed on an Orbitrap Fusion Lumos Tribrid using a multinotch SPS-MS³ TMT method^75^. Full MS scans (400–1,400 m/z) were acquired in the Orbitrap with 2 min dynamic exclusion. MS² (turbo) scans were recorded in the ion trap with CID 35% and maximum injection times up to 450 ms. SPS-MS³ spectra were collected in the Orbitrap (100–1,000 m/z) after HCD 55%, 50,000 resolution, and injection times up to 600 ms.

#### MS Data Analysis

Spectra were searched with SEQUEST (v28 rev12) against a Mus musculus UniProt (03/2021) database plus contaminants. Precursor and fragment tolerances were 20 ppm and 0.9 Da, respectively; up to two missed cleavages were allowed. Dynamic modifications: Met +15.9949 Da. Static modifications: Cys +57.0215 Da (IAA) and TMTpro +229.1629 Da on peptide N-termini and Lys. Target–decoy filtering with linear discriminant analysis set peptide-spectrum match and protein FDR at 1%. MS data were recalibrated post-search and re-searched. Quantification used SPS-MS^3^ spectra with ≥70% isolation specificity and summed reporter signal-to-noise ≥200 across TMT channels. Detailed quantification procedures and additional LC–MS settings were as described^76,77^.

### Retinal Methods

#### Staining of Zebrafish retina

Immunofluorescence was performed as previously described^78^. In brief, larvae were fixed in 4% paraformaldehyde in phosphate buffer at room temperature for 2 hours, then embedded in tissue freezing medium (Electron Microscopy Sciences, Hatfield, PA, USA) and cryo-sectioned. To bleach pigmentation, sections were treated with 10% hydrogen peroxide in PBS at 65 °C for 15 minutes. Unspecific binding was blocked using PBDT buffer (PBS containing 0.5% Triton X-100, 1% BSA, and 1% DMSO) supplemented with 10% goat serum for 2 hr at room temperature. Sections were incubated overnight at 4 °C with the primary antibody against ubiquitin (FK2, 1:100 in PBDT). After four washes in PBDT, sections were incubated with goat anti-mouse Alexa Fluor 594-conjugated secondary antibody (1:400) for 2.5 hr at room temperature. Membranes of the outer segments and nuclei were counterstained using Vybrant DiO (1:200) and DAPI (5 µg/mL), respectively, diluted in PDT buffer (PBS containing 0.1% Triton X-100 and 1% DMSO) for 20 minutes at room temperature. Imaging was performed using a Leica Stellaris 5 upright confocal microscope.

#### AAV packaging and subretinal injection

Adeno-associated virus serotype 5 (AAV5) were produced by Vector Builder by co-transfecting the shuttle plasmids encoding IMPG2 or IMPG2[cK0] with Rep-cap plasmid and helper plasmid encoding adenovirus genes (E4, E2A and VA) into HEK293T packaging cells to promote AAV replication. Viral particles were harvested from cell lysates, concentrated by PEG precipitation and further purified and concentrated by cesium chloride gradient ultracentrifugation. AAV titers were measured by qPCR. The final formulations of the viral vectors were 6.1 × 10^13^ genome copies (GC) per mL (IMPG2) and 5.3 × 10^13^ GC/mL (IMPG2[cK0]) in PBS supplemented with 200 mM NaCl and 0.001% pluronic F-68.

Subretinal injections of 1-2 µL AAV5 viral vectors encoding IMPG2 variants were performed on neonatal wildtype CD-1 mice at postnatal day 2, as previously described^79^. Both male and female wildtype littermate mice were randomly assigned to experimental groups.

Mice were euthanized three weeks post-injection, and retinal sections were processed for immunofluorescence. Images were acquired on a LSM900 Zeiss microscope equipped with Plan Apochromat 40X/1.4 DIC Oil objective in Z stacks of 12 to 15 optical sections captured at 1µm interval.

#### Staining and imaging of retinal tissue sections

For P8 animals, eyes were slit open with a needle. Mouse eyes were enucleated, and fixed in 4% (w/v) paraformaldehyde in PBS for 30 minutes. After cornea, lens and vitreous humor removal, the eye cups were further fixed in the same paraformaldehyde solution for another 30 minutes. Eyecups were washed in PBS then cryoprotected in 30% sucrose for 1 hour. Eyecups were embedded in Optimal Cutting Temperature (OCT) compound medium (4583; Tissue-Tek) in a mold, frozen on dry ice and stored at −80 °C. 12 μm-sections were cut on a cryostat (CM 1860; Leica). Retinal sections were blocked in wash buffer (50 mM Tris, pH7.4, 100 mM NaCl, 0.1% Triton X-100) supplemented with 3% normal donkey serum and 0.1% BSA for 1 hr at room temperature and incubated overnight at 4 °C with primary antibodies diluted in wash buffer with blocking reagents. Sections were then washed three times with wash buffer and incubated with secondary antibodies diluted in wash buffer for 2 hr at room temperature. After three additional washes, DNA was counterstained using Hoescht 33342 (diluted 1:10,000 in wash buffer) and washed twice more with PBS before mounted onto slides with Fluormount G.

Staining with the GST-TUBE/GST proteins was conducted by using the GST fusion (diluted to 10 µg/mL in wash buffer) in lieu of primary antibodies and using Alexa Fluor 488-conjugated rabbit polyclonal anti-GST antibodies (diluted 1:500 in wash buffer) in lieu of secondary antibodies.

Retinal sections were imaged on LSM700 (for **Figs.2, 4C, 4E, S1A, S1C-D, S2, S3**) or LSM900 (for **Figs. 1, 4G, 7**, **S1B, S5**) (Zeiss) microscopes with a Plan-Apochromat 40X/1.4 DIC Oil objective at 1 µm interval in Z stacks of 12-15 planes.

#### Image analysis of retinal tissue sections

Confocal retinal images acquired on Zeiss LSM700 and LSM900 microscopes were imported into ImageJ/Fiji (National Institutes of Health) for analysis. To quantify photoreceptor signals in the outer and inner retinal segments, maximum intensity projections were generated from 6 to 11 Z-sections with uninterrupted morphology. Signals were measured in three circular regions of interest (ROIs) distributed in the relevant areas across the entire field of view (4 µm diameter for P8; 20 µm diameter for P11 and older). Integrated density values from these ROIs were used to calculate the ratio of outer to inner segment signals, with results plotted as individual data points using PlotsofData.

In **Fig. 7C**, IMP2^m^ intensity values in the OS and IS were background-corrected using corresponding IMPG2^m^ intensity values from the uninjected areas of the same retina and OS signals were then normalized against IS signals.

## BIBLIOGRAPHY

1. Forsythe, E. & Beales, P. L. Bardet-Biedl Syndrome. in GeneReviews® (eds. Adam, M. P. et al.) (University of Washington, Seattle, Seattle (WA), 1993).

2. Nachury, M. V. et al. A core complex of BBS proteins cooperates with the GTPase Rab8 to promote ciliary membrane biogenesis. Cell 129, 1201–1213 (2007).

3. Nachury, M. V. The molecular machines that traffic signaling receptors into and out of cilia. Curr. Opin. Cell Biol. 51, 124–131 (2018).

4. Tian, X., Zhao, H. & Zhou, J. Organization, functions, and mechanisms of the BBSome in development, ciliopathies, and beyond. Elife 12, e87623 (2023).

5. Wingfield, J. L., Lechtreck, K.-F. & Lorentzen, E. Trafficking of ciliary membrane proteins by the intraflagellar transport/BBSome machinery. Essays Biochem. 62, 753–763 (2018).

6. Nachury, M. V. & Mick, D. U. Establishing and regulating the composition of cilia for signal transduction. Nat. Rev. Mol. Cell Biol. 20, 389–405 (2019).

7. Lacey, S. E. & Pigino, G. The intraflagellar transport cycle. Nat Rev Mol Cell Biol (2024) doi:10.1038/s41580-024-00797-x.

8. Yau, R. & Rape, M. The increasing complexity of the ubiquitin code. Nature Cell Biology 18, 579–586 (2016).

9. Swatek, K. N. & Komander, D. Ubiquitin modifications. Cell Research 26, 399–422 (2016).

10. Shinde, S. R., Nager, A. R. & Nachury, M. V. Ubiquitin chains earmark GPCRs for BBSome-mediated removal from cilia. J Cell Biol 219, e202003020 (2020).

11. Desai, P. B., Stuck, M. W., Lv, B. & Pazour, G. J. Ubiquitin links smoothened to intraflagellar transport to regulate Hedgehog signaling. J Cell Biol 219, e201912104 (2020).

12. Piper, R. C., Dikic, I. & Lukacs, G. L. Ubiquitin-Dependent Sorting in Endocytosis. Cold Spring Harbor Perspectives in Biology 6, a016808–a016808 (2014).

13. May, E. A., Sroka, T. J. & Mick, D. U. Phosphorylation and Ubiquitylation Regulate Protein Trafficking, Signaling, and the Biogenesis of Primary Cilia. Front Cell Dev Biol 9, 664279 (2021).

14. Shinde, S. R. et al. The ancestral ESCRT protein TOM1L2 selects ubiquitinated cargoes for retrieval from cilia. Developmental Cell 58, 1–17 (2023).

15. Datta, P. et al. Accumulation of non-outer segment proteins in the outer segment underlies photoreceptor degeneration in Bardet-Biedl syndrome. Proc Natl Acad Sci U S A 112, E4400–9 (2015).

16. Baker, S. A. et al. The outer segment serves as a default destination for the trafficking of membrane proteins in photoreceptors. The Journal of Cell Biology 183, 485–498 (2008).

17. Seo, S. & Datta, P. Photoreceptor outer segment as a sink for membrane proteins: hypothesis and implications in retinal ciliopathies. Hum Mol Genet 26, R75–R82 (2017).

18. Garcia-Gonzalo, F. R. & Reiter, J. F. Open Sesame: how transition fibers and the transition zone control ciliary composition. Cold Spring Harb Perspect Biol 9, a028134 (2017).

19. Gonçalves, J. & Pelletier, L. The Ciliary Transition Zone: Finding the Pieces and Assembling the Gate. Molecules and cells 40, 243–253 (2017).

20. Datta, P., Cribbs, J. T. & Seo, S. Differential requirement of NPHP1 for compartmentalized protein localization during photoreceptor outer segment development and maintenance. PLoS One 16, e0246358 (2021).

21. Datta, P., Hendrickson, B., Brendalen, S., Ruffcorn, A. & Seo, S. The myosin-tail homology domain of centrosomal protein 290 is essential for protein confinement between the inner and outer segments in photoreceptors. J. Biol. Chem. jbc.RA119.009712 (2019) doi:10.1074/jbc.RA119.009712.

22. Truong, H. M. et al. The tectonic complex regulates membrane protein composition in the photoreceptor cilium. Nat Commun 14, 5671 (2023).

23. Nishimura, D. Y. et al. Bbs2-null mice have neurosensory deficits, a defect in social dominance, and retinopathy associated with mislocalization of rhodopsin. Proceedings of the National Academy of Sciences of the United States of America 101, 16588–16593 (2004).

24. Davis, R. E. et al. A knockin mouse model of the Bardet-Biedl syndrome 1 M390R mutation has cilia defects, ventriculomegaly, retinopathy, and obesity. Proc Natl Acad Sci U S A 104, 19422–19427 (2007).

25. Mykytyn, K. et al. Bardet-Biedl syndrome type 4 (BBS4)-null mice implicate Bbs4 in flagella formation but not global cilia assembly. Proceedings of the National Academy of Sciences of the United States of America 101, 8664–8669 (2004).

26. Dilan, T. L. et al. Bardet-Biedl syndrome-8 (BBS8) protein is crucial for the development of outer segments in photoreceptor neurons. Hum Mol Genet 27, 283–294 (2018).

27. Pearring, J. N., Salinas, R. Y., Baker, S. A. & Arshavsky, V. Y. Protein sorting, targeting and trafficking in photoreceptor cells. Progress in retinal and eye research 36, 24–51 (2013).

28. Sedmak, T. & Wolfrum, U. Intraflagellar transport molecules in ciliary and nonciliary cells of the retina. The Journal of Cell Biology 189, 171–186 (2010).

29. Masek, M. et al. Loss of the Bardet-Biedl protein Bbs1 alters photoreceptor outer segment protein and lipid composition. Nat Commun 13, 1282 (2022).

30. Hjerpe, R. et al. Efficient protection and isolation of ubiquitylated proteins using tandem ubiquitin-binding entities. EMBO Rep. 10, 1250–1258 (2009).

31. Sims, J. J. et al. Polyubiquitin-sensor proteins reveal localization and linkage-type dependence of cellular ubiquitin signaling. Nat Methods 9, 303–309 (2012).

32. Skiba, N. P. et al. TMEM67, TMEM237, and Embigin in Complex With Monocarboxylate Transporter MCT1 Are Unique Components of the Photoreceptor Outer Segment Plasma Membrane. Molecular & Cellular Proteomics 20, 100088 (2021).

33. Hospenthal, M. K., Mevissen, T. E. T. & Komander, D. Deubiquitinase-based analysis of ubiquitin chain architecture using Ubiquitin Chain Restriction (UbiCRest). Nat Protoc 10, 349–361 (2015).

34. Liu, G. et al. Mechanism of adrenergic CaV1.2 stimulation revealed by proximity proteomics. Nature 577, 695–700 (2020).

35. Li, J. et al. TMTpro-18plex: The Expanded and Complete Set of TMTpro Reagents for Sample Multiplexing. J Proteome Res 20, 2964–2972 (2021).

36. Li, J. et al. TMTpro reagents: a set of isobaric labeling mass tags enables simultaneous proteome-wide measurements across 16 samples. Nat. Methods 17, 399–404 (2020).

37. Curtis, L. B. et al. Syntaxin 3b is a t-SNARE specific for ribbon synapses of the retina. Journal of Comparative Neurology 510, 550–559 (2008).

38. Mazelova, J., Ransom, N., Astuto-Gribble, L., Wilson, M. C. & Deretic, D. Syntaxin 3 and SNAP-25 pairing, regulated by omega-3 docosahexaenoic acid, controls the delivery of rhodopsin for the biogenesis of cilia-derived sensory organelles, the rod outer segments. Journal of Cell Science 122, 2003–2013 (2009).

39. Chuang, J.-Z., Zhao, Y. & Sung, C.-H. SARA-regulated vesicular targeting underlies formation of the light-sensing organelle in mammalian rods. Cell 130, 535–547 (2007).

40. Hsu, Y. et al. BBSome function is required for both the morphogenesis and maintenance of the photoreceptor outer segment. PLoS Genet. 13, e1007057 (2017).

41. Giovannone, A. J., Reales, E., Bhattaram, P., Fraile-Ramos, A. & Weimbs, T. Monoubiquitination of syntaxin 3 leads to retrieval from the basolateral plasma membrane and facilitates cargo recruitment to exosomes. Molecular Biology of the Cell 28, 2843–2853 (2017).

42. Daniele, L. L., Adams, R. H., Durante, D. E., Pugh, E. N. & Philp, N. J. Novel distribution of junctional adhesion molecule-C in the neural retina and retinal pigment epithelium. J Comp Neurol 505, 166–176 (2007).

43. Kwok, M. C. M., Holopainen, J. M., Molday, L. L., Foster, L. J. & Molday, R. S. Proteomics of Photoreceptor Outer Segments Identifies a Subset of SNARE and Rab Proteins Implicated in Membrane Vesicle Trafficking and Fusion. Molecular & Cellular Proteomics 7, 1053–1066 (2008).

44. Spencer, W. J. et al. The WAVE complex drives the morphogenesis of the photoreceptor outer segment cilium. Proc Natl Acad Sci U S A 120, e2215011120 (2023).

45. Ishikawa, M., Sawada, Y. & Yoshitomi, T. Structure and function of the interphotoreceptor matrix surrounding retinal photoreceptor cells. Experimental Eye Research 133, 3–18 (2015).

46. Mitchell, B., Coulter, C., Geldenhuys, W. J., Rhodes, S. & Salido, E. M. Interphotoreceptor matrix proteoglycans IMPG1 and IMPG2 proteolyze in the SEA domain and reveal localization mutual dependency. Sci Rep 12, 15535 (2022).

47. Salido, E. M. & Ramamurthy, V. Proteoglycan IMPG2 Shapes the Interphotoreceptor Matrix and Modulates Vision. J. Neurosci. 40, 4059–4072 (2020).

48. Hsu, Y., Seo, S. & Sheffield, V. C. Photoreceptor cilia, in contrast to primary cilia, grant entry to a partially assembled BBSome. Hum Mol Genet (2021) doi:10.1093/hmg/ddaa284.

49. Zhang, M., Berk, J. M., Mehrtash, A. B., Kanyo, J. & Hochstrasser, M. A versatile new tool derived from a bacterial deubiquitylase to detect and purify ubiquitylated substrates and their interacting proteins. PLoS Biol 20, e3001501 (2022).

50. Lv, B., Stuck, M. W., Desai, P. B., Cabrera, O. A. & Pazour, G. J. E3 ubiquitin ligase Wwp1 regulates ciliary dynamics of the Hedgehog receptor Smoothened. J Cell Biol 220, e202010177 (2021).

51. Acharya, S. et al. SPACRCAN, a Novel Human Interphotoreceptor Matrix Hyaluronan-binding Proteoglycan Synthesized by Photoreceptors and Pinealocytes. Journal of Biological Chemistry 275, 6945–6955 (2000).

52. Hollyfield, J. G., Rayborn, M. E., Midura, R. J., Shadrach, K. G. & Acharya, S. Chondroitin Sulfate Proteoglycan Core Proteins in the Interphotoreceptor Matrix: A Comparative Study using Biochemical and Immunohistochemical Analysis. Experimental Eye Research 69, 311–322 (1999).

53. Park, K. & Leroux, M. R. Composition, organization and mechanisms of the transition zone, a gate for the cilium. EMBO reports 23, e55420 (2022).

54. Shi, X. et al. Super-resolution microscopy reveals that disruption of ciliary transition-zone architecture causes Joubert syndrome. Nat. Cell Biol. 19, 1178–1188 (2017).

55. Ye, F., Nager, A. R. & Nachury, M. V. BBSome trains remove activated GPCRs from cilia by enabling passage through the transition zone. J. Cell Biol. 217, 1847–1868 (2018).

56. Barbelanne, M., Hossain, D., Chan, D. P., Peränen, J. & Tsang, W. Y. Nephrocystin proteins NPHP5 and Cep290 regulate BBSome integrity, ciliary trafficking and cargo delivery. Human Molecular Genetics 24, 2185–2200 (2015).

57. Zhang, Y. et al. BBS mutations modify phenotypic expression of CEP290-related ciliopathies. Human Molecular Genetics 23, 40–51 (2014).

58. Yee, L. E. et al. Conserved Genetic Interactions between Ciliopathy Complexes Cooperatively Support Ciliogenesis and Ciliary Signaling. PLoS Genet 11, e1005627 (2015).

59. Goetz, S. C., Bangs, F., Barrington, C. L., Katsanis, N. & Anderson, K. V. The Meckel syndrome-associated protein MKS1 functionally interacts with components of the BBSome and IFT complexes to mediate ciliary trafficking and hedgehog signaling. PLoS ONE 12, e0173399 (2017).

60. Bentley-Ford, M. R. et al. Evolutionarily conserved genetic interactions between nphp-4 and bbs-5 mutations exacerbate ciliopathy phenotypes. Genetics 220, iyab209 (2022).

61. Zulliger, R. et al. SNAREs Interact with Retinal Degeneration Slow and Rod Outer Segment Membrane Protein-1 during Conventional and Unconventional Outer Segment Targeting. PLoS One 10, e0138508 (2015).

62. Kakakhel, M. et al. Syntaxin 3 is essential for photoreceptor outer segment protein trafficking and survival. Proc Natl Acad Sci USA 117, 20615–20624 (2020).

63. Janecke, A. R. et al. Pathogenic STX3 variants affecting the retinal and intestinal transcripts cause an early-onset severe retinal dystrophy in microvillus inclusion disease subjects. Hum Genet 140, 1143–1156 (2021).

64. Bandah-Rozenfeld, D. et al. Mutations in IMPG2, encoding interphotoreceptor matrix proteoglycan 2, cause autosomal-recessive retinitis pigmentosa. Am J Hum Genet 87, 199–208 (2010).

65. Mockel, A. et al. Pharmacological modulation of the retinal unfolded protein response in Bardet-Biedl syndrome reduces apoptosis and preserves light detection ability. Journal of Biological Chemistry 287, 37483–37494 (2012).

66. Tumbarello, D. A. et al. Autophagy receptors link myosin VI to autophagosomes to mediate Tom1-dependent autophagosome maturation and fusion with the lysosome. Nature Cell Biology 14, 1024–1035 (2012).

67. Perez-Riverol, Y. et al. The PRIDE database at 20 years: 2025 update. Nucleic Acids Res gkae1011 (2024) doi:10.1093/nar/gkae1011.

68. Jin, H. et al. The conserved Bardet-Biedl syndrome proteins assemble a coat that traffics membrane proteins to cilia. Cell 141, 1208–1219 (2010).

69. Aleström, P. et al. Zebrafish: Housing and husbandry recommendations. Lab Anim 54, 213–224 (2020).

70. Sims, J. J. et al. Polyubiquitin-sensor proteins reveal localization and linkage-type dependence of cellular ubiquitin signaling. Nat Methods 9, 303–309 (2012).

71. Hjerpe, R. et al. Efficient protection and isolation of ubiquitylated proteins using tandem ubiquitin-binding entities. EMBO reports 10, 1250–1258 (2009).

72. Liew, G. M. et al. The intraflagellar transport protein IFT27 promotes BBSome exit from cilia through the GTPase ARL6/BBS3. Dev Cell 31, 265–278 (2014).

73. Postma, M. & Goedhart, J. PlotsOfData-A web app for visualizing data together with their summaries. PLoS Biol 17, e3000202 (2019).

74. Hughes, C. S. et al. Single-pot, solid-phase-enhanced sample preparation for proteomics experiments. Nat Protoc 14, 68–85 (2019).

75. McAlister, G. C. et al. MultiNotch MS3 enables accurate, sensitive, and multiplexed detection of differential expression across cancer cell line proteomes. Anal Chem 86, 7150–7158 (2014).

76. Huttlin, E. L. et al. A tissue-specific atlas of mouse protein phosphorylation and expression. Cell 143, 1174–1189 (2010).

77. Paulo, J. A. et al. Quantitative mass spectrometry-based multiplexing compares the abundance of 5000 S. cerevisiae proteins across 10 carbon sources. J Proteomics 148, 85–93 (2016).

78. Masek, M. et al. Chapter 7 - Studying the morphology, composition and function of the photoreceptor primary cilium in zebrafish. in Methods in Cell Biology (eds. Bravo-San Pedro, J. M. & Galluzzi, L.) vol. 175 97–128 (Academic Press, 2023).

79. Wert, K. J., Skeie, J. M., Davis, R. J., Tsang, S. H. & Mahajan, V. B. Subretinal Injection of Gene Therapy Vectors and Stem Cells in the Perinatal Mouse Eye. J Vis Exp 4286 (2012) doi:10.3791/4286.

